# Dosing and Delivery of Bacteriophage Therapy In a Murine Wound Infection Model

**DOI:** 10.1101/2024.05.07.593005

**Authors:** Yung-Hao Lin, Tejas Dharmaraj, Qingquan Chen, Arne Echterhof, Robert Manasherob, Lucy J. Zhang, Zhiwei Li, Cas de Leeuw, Nana A. Peterson, Tony H. W. Chang, Whitney Stannard, Maryam Hajfathalian, Aviv Hargil, Hunter A. Martinez, Julie Pourtois, Francis G. Blankenberg, Derek Amanatullah, Ovijit Chaudhuri, Paul L. Bollyky

## Abstract

Lytic bacteriophages, viruses that lyse (kill) bacteria, hold great promise for treating infections, including wound infections caused by antimicrobial-resistant *Pseudomonas aeruginosa.* However, dosing and delivery strategies for phage therapy remain underdeveloped. In a mouse wound infection model, we investigated the impact of administration route, dose, and frequency on the efficacy of phage therapy. We find that topical but not systemic delivery is effective in this model. *In vitro* and *in vivo* data supported the use of high doses of phage. Repeated dosing achieves the highest eradication rates *in vivo.* Building on these insights, we developed “HydroPhage”, a hyaluronan-based hydrogel system that uses dynamic covalent crosslinking to deliver high-titre phages over one week, a substantial improvement over existing burst-release systems. We conclude that hydrogel-based sustained phage delivery offers a practical, efficacious, and well-tolerated option for topical phage application.

## Introduction

Chronic wounds are frequently complicated by antimicrobial-resistant (AMR) infections^1^, with *Pseudomonas aeruginosa* implicated in 25-50% of chronically infected wounds^2–5^. Reducing the bacterial burden in chronic wounds is an essential step in wound care^6^. Notably, *P. aeruginosa* strains in chronic wounds are frequently multidrug-resistant (MDR)^7–14^. MDR *P. aeruginosa* is recognized as a serious threat by both the Centers for Disease Control and Prevention (CDC) and the World Health Organization (WHO)^15,16^, with few conventional antibiotics in development^17^. This underscores the urgent need to pursue alternative therapies, including phage therapy (PT).

Bacteriophages (phages), viruses that infect and kill bacteria^19^, represent a promising alternative for treating bacterial infections. PT offers several advantages, including targeted activity against bacterial pathogens^18^, biofilm disruption^19^, and favorable safety profiles^20–23^. Numerous studies have demonstrated the success of PT in treating MDR *P. aeruginosa* infections^24–26^. However, despite advances in phage pharmacology^27,28^, there remains a lack of preclinical and clinical evidence to guide PT dosing and administration for infected wounds. Consequently, the use of PT for wound infections is still in its clinical infancy.

For many indications, PT in humans is typically administered intravenously^37^, although local delivery is recommended whenever possible^38^. Clinical trials, often based on anecdotal or theoretical foundations, have employed repeated topical applications of phages, either as sprays or gauze soaked with phage suspensions, to treat wound infections^29,30^. While these methods have shown promise, there is a lack of preclinical or clinical data supporting the concept topical treatment is more effective than systemic treatment. Moreover, repeated topical dosing necessitates inpatient care or frequent patient follow-up, highlighting the need for advanced wound dressings that incorporate phages.

Hydrogel-based dressings, which provide a moist environment conducive to healing, have emerged as a promising solution. However, existing burst-release hydrogels lack controlled, sustained phage release, which is critical for maintaining therapeutic levels over time and effectively mimicking repeated dosing in a single application. Thus, there is a pressing need for advanced approaches that enable controlled phage release, optimize therapeutic efficacy, and reduce the frequency of reapplications.

Here, we employed an established mouse model^31–33^–modified to feature extended wound infection and tailored for PT studies–to assess the effectiveness of both systemic and topical PT in reducing bacterial burdens in wounds. In parallel, we designed a phage-delivering hydrogel capable of sustained phage release (**Fig. 1a**).

**Fig. 1.**
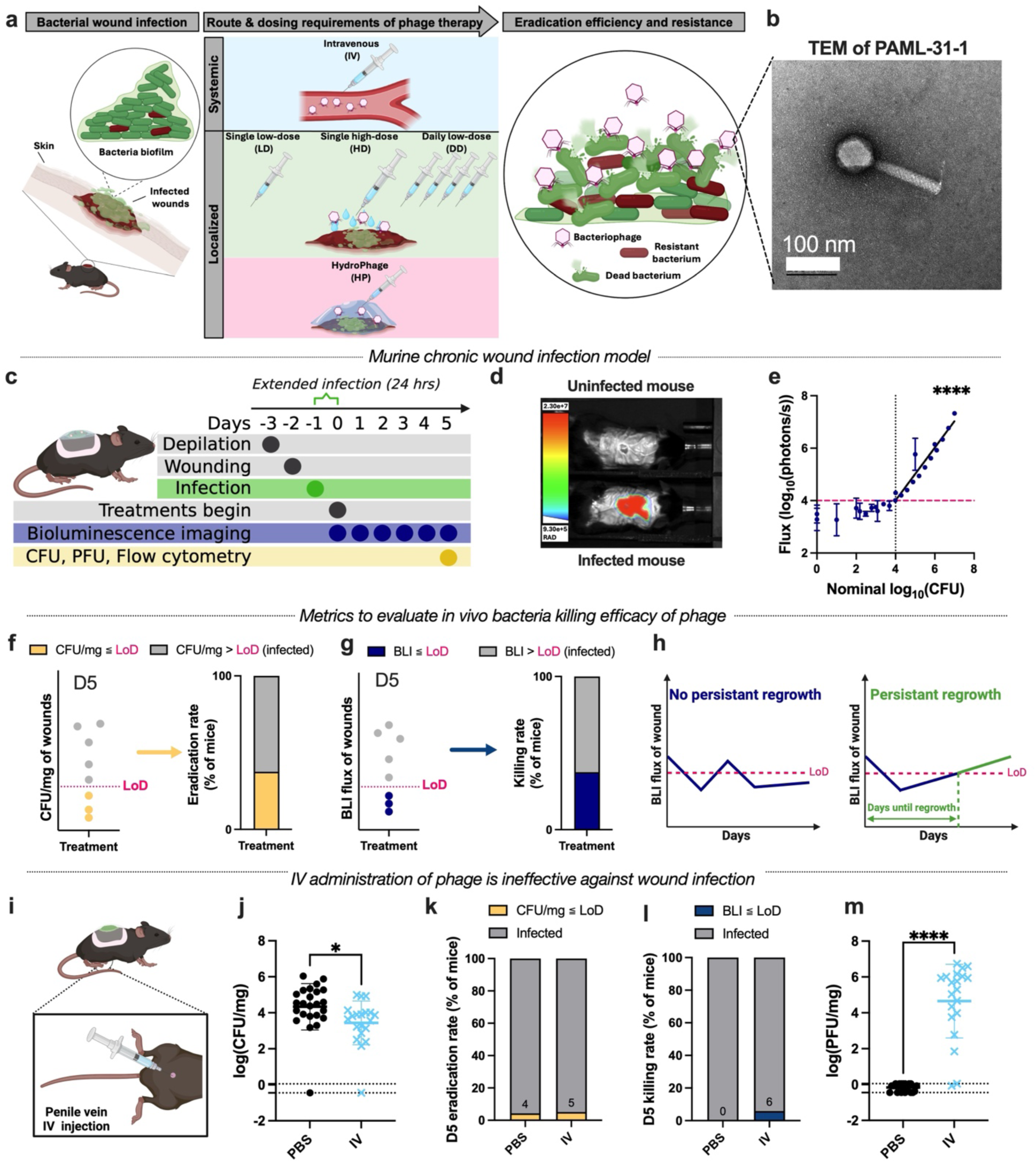
Murine model shows intravenous (IV) phage therapy (PT) is ineffective against *P. aeruginosa* wound infection. **a**, illustration of the research summary. **b**, PAML-31-1 phage visualized using transmission electron microscopy (TEM). Scale bar = 100 nm. **c**,**d**,**e**, Development of murine wound infection model. Illustration of the timeline of murine wound model (**c**); the representative bioluminescence imaging (BLI) image of uninfected (top) and infected (bottom) mice, where color bar indicates the flux intensity (**d**). The infected mouse image was selected from PBS-treated group on D2 post-infection; bioluminescent signal and nominal CFU correlation above above 1×10^4^ CFU for PAO1-Xen41 *in vitro* (Pearson r = 0.953, P < 0.001; R^2^ = 0.889) (**e**); **f**,**g**,**h**, Metrics used to evaluate the *in vivo* bacterial killing efficacy of phage treatments. The eradication rate defined as the percentage of mice with wound CFU/mg below the limit of detection (LoD) on Day 5 post-treatment (**f**); The killing rate defined as the percentage of mice with wounds having bioluminescent flux signal below the LoDon Day 5 post-treatment (**g**); the persistent growth rate defined as the percentage of mice with wound bioluminescent flux signal remained higher than LoD (**h**). **i**,**j**,**k**,**l**,**m**, Evaluation of *in vivo* efficacy of IV phage treatment. Illustration demonstrates the penile vein injection for IV phage treatment (1 x 10^11^ PFU/mL, 40 µL, 4 x 10^9^ PFU total) (**i**); bacterial burden (CFU/mg) recovered from harvested mice wound tissue (**j**); the eradiation rate (**k**) for PBS and IV groups, n=24 and 20 mice, n=5 and 2 independent experiments, respectively. The killing rate (**l**) for PBS and IV groups, n=17 and 17 mice, n=4 and 2 independent experiments, respectively. The phage counts (PFU/mg) recovered from harvested mice wound tissue (**m**). Schematics in (**a**), (**c**) and (**i**) were generated with BioRender. Data are presented as mean ± S.D. Linear fit and Pearson r for (**e**). Unpaired two-tailed Welch’s *t*-test for (**j**) and (**m**). (* *P* ≤ 0.05, **** *P* ≤ 0.0001).

## Results

### A murine model of extended infection

We adapted the delayed inoculation wound infection model to study PT ^31–33^. This model involves depilation on Day -3, wounding on Day -2, and inoculation of the wound with PAO1-Xen41 on Day -1 for 24 hrs to establish infections. PAO1-Xen41 is a luminescent derivative of the common *P. aeruginosa* lab strain PAO1^34^. Inoculations were performed under an occlusive dressing (Tegaderm) to prevent contamination. By allowing a provisional wound matrix to form before inoculation, this model enables stable *P. aeruginosa* colonization without the need for foreign material or immune suppression. PT was then introduced to established infections one day after inoculation, on Day 0. An overview of this protocol is shown in **Figure 1c**.

We selected a phage with high activity against PAO1-Xen41 by evaluating virulent antipseudomonal phages PAML-31-1, LPS-5, Luz24, and OMKO1, using planktonic suppression assays (PSA) and semi-solid substrate tests –efficiency of plaquing (EOP) **(Extended Data Fig. 1a)**. Among them, PAML-31-1 **(Fig. 1b)**, a Pbunavirus which targets pseudomonal oligosaccharide antigen/lipopolysaccharide (OSA/LPS), demonstrated the strongest suppression of PAO1-Xen41, with the highest “Suppression Index” (SI%)^35^ –the percentage of growth inhibition caused by phage treatment within the first 48 hours of exposure **(Extended Data Fig. 1a-k)** in the PSA and the highest EOP against PAO1-Xen41 **(Extended Data Fig. 1l)**. Therefore, PAML-31-1 was selected for further studies.

We then identified that an inoculum of 5 x 10^5^ colony-forming units (CFU)/wound of PAO1-Xen41 was the minimum required to establish a 100% infection rate in mice, by testing varying bacterial loads of PAO1-Xen41 **(Extended Data Fig. 2a)**. Infection rate was defined as the percentage of mice that had CFU counts above the limit of detection (LoD) at the endpoint of the study, Day 6 post-infection. In addition to standard culture-based methods to enumerate CFU at the study endpoint on Day 5 post-infection, we tracked the progress of the infections daily using bioluminescence imaging (BLI) **(Fig. 1d)**. CFU were closely correlated with luminescent flux **(Fig. 1e and Extended Data Fig. 2b)**. We observed that infections caused pain, irritation, and redness at the wound site. Still, mice showed minimal systemic symptoms, with 100% survival (**Extended Data Fig. 2c**) and slight weight loss (**Extended Data Fig. 2d**).

This robust preclinical model allowed for assessing PT efficacy using CFU enumeration at the study endpoint with simultaneous real-time BLI monitoring to corroborate the results. Our primary endpoints were 1) reduction of bacterial burdens in wounds (quantified as log reduction in CFU), 2) endpoint eradication rate (percentage of mice with wound CFU/mg below LoD) and killing rate (percentage of mice with wound BLI below LoD), and 3) persistent regrowth rate (percentage of mice with wound BLI flux irreversibly above LoD over time) **(Fig. 1f–h)**. Secondary endpoints included the quantification of phage in the wound tissue at the study endpoint as plaque-forming units (PFU), the cellular composition of the wound immune microenvironment in response to treatment, and the emergence of resistance to PT.

### Systemic PT does not reduce the bacterial burden in infected murine wounds

We first investigated the efficacy of systemic PT in reducing bacterial burdens in the wound bed. As per the protocol in **Figure 1b**, mice were wounded, inoculated, and treated the following day with either topical PBS or PAML-31-1 administered intravenously (IV) via the dorsal penile vein (1 x 10^11^ PFU/mL, 40 µL, 4 x 10^9^ PFU total) **(Fig. 1i).** Culture-based assays showed minor mean log reduction (0.90) in CFU/mg tissue at the study endpoint **(Fig. 1j)**. PBS and IV phage led to only 5-6% eradication rate **(Fig. 1k)**. The killing rate derived from BLI imaging corroborated with culture-based result, showing 0% for PBS and 6% for IV phage treatment **(Fig. 1l)**. Repeated IV injections were avoided due to low cumulative accuracy and potential damage to soft tissues^36^. Thus, as an alternative approach to systemic and repeated PT, we performed intraperitoneally (IP) as four divided doses, each administered daily (IP*4) (2.5 x 10^9^ PFU/mL, 400 µL/dose, 4 x 10^9^ PFU total). Similary, we observed no significant mean log reduction (0.34) in CFU/mg **(Extended Data Fig.3a)** and only 7% eradication rate **(Extended Data Fig. 3b).** These results confirmed existing anecdotal knowledge that systemic PT is ineffective for reducing bacterial burdens in wound infections, despite substantial presence of phage in homogenized wound tissue at the study endpoint **(Fig. 1m and Extended Data Fig. 3c).**

### Localized free phage delivery reduces the bacterial burden in infected murine wounds

To evaluate the therapeutic potential of localized free phage delivery, we administered high-titer phage treatments to infected murine wounds **(Fig. 2a)**. Mice received either a single high-dose (HD, 1 x 10^11^ PFU/mL, 40 µL, 4 x 10^9^ PFU total), a single low-dose treatment (LD, 2.5 x 10^10^ PFU/mL, 40 µL, 1 x 10^9^ PFU total), or a daily dose for 4 days (DD, 2.5 x 10^10^ PFU/mL/day, 40 µL, 4 x 10^9^ PFU total), which is equivalent to low-dose administered daily or high-dose given as four divided doses) **(Fig. 2b)**. High phage titers were confirmed in wound homogenates at the study endpoint across all treatment groups (**Fig. 2c**).

**Fig. 2.**
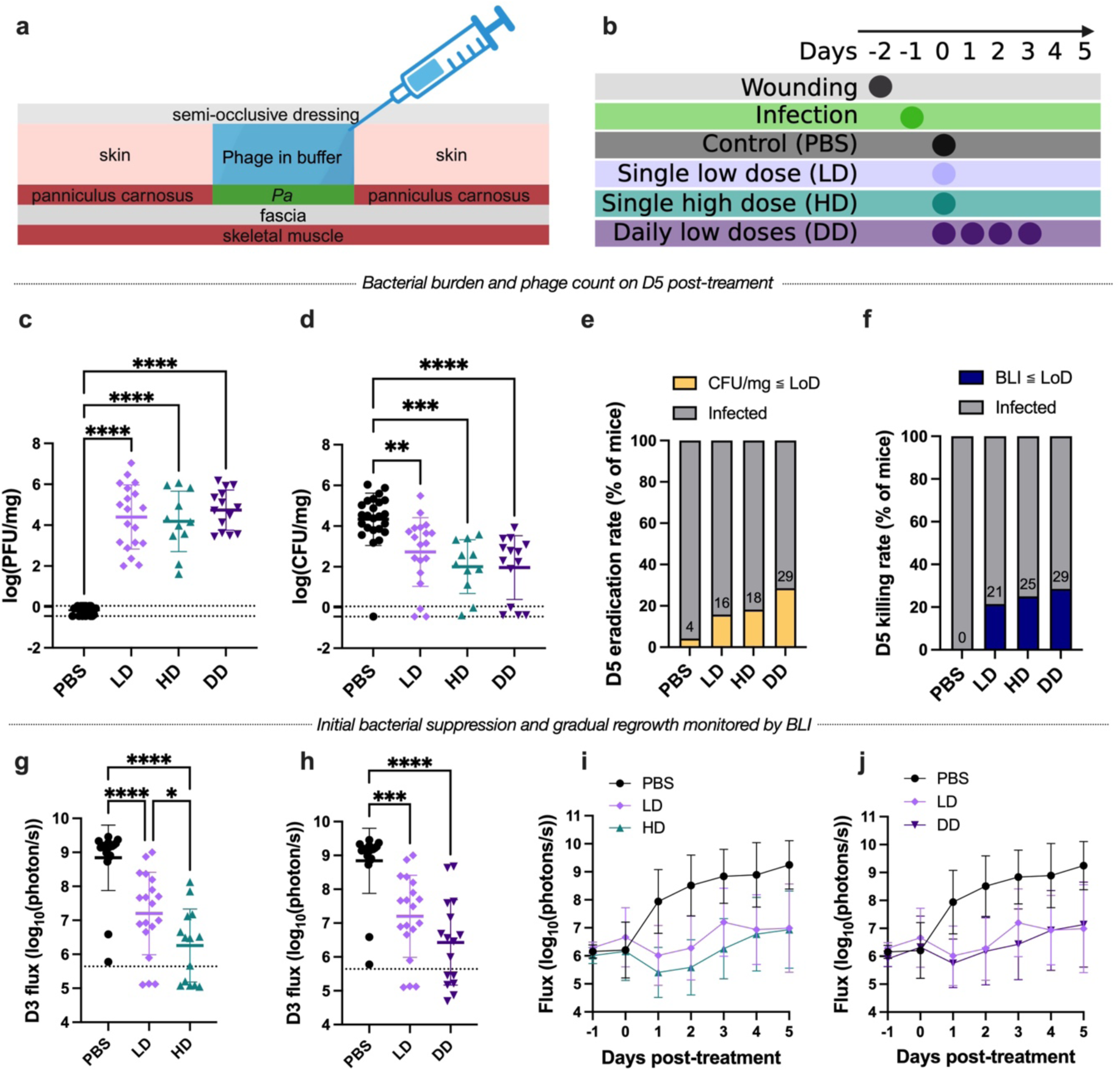
Local phage delivery reduces bacterial burden. **a**, Illustration of how mice receive liquid phage treatments by topical injection under a Tegaderm semi-occlusive dressing. **b,** Illustration of the murine wound study to compare the efficacy of phage therapy (PT) with different dosing concentration and frequency. **c**,**d**, the phage counts (PFU/mg) (**c**) and bacterial burden (CFU/mg) (**d**) recovered from harvested mice wound tissue, n=24, 19, 11, 14 mice and n=5, 2, 2, 2 independent experiments for PBS, LD, HD and DD groups, respectively. **e**, The eradication rate (CFU-based) achieved by treatments on Day 5 post-treatment, n=24, 19, 11, 14 mice and n=5, 2, 2, 2 independent experiments for PBS, LD, HD, and DD groups, respectively. **f**, The killing rate (BLI-based) achieved by treatments on Day 5 post-treatment, n=17, 14, 12, 14 mice and n=4, 2, 2, 2 independent experiments for PBS, LD, HD, and DD groups, respectively. **g**,**h**, The bioluminescent flux signal on Day 3 post-treatment comparing total dose (PBS, LD, and HD) (**g**) and frequency of dosing (PBS, LD, and DD) (**h**), n=19, 19, 14, 16 for PBS, LD, HD, and DD groups, respectively; from 4, 2, 2, 2 independent experiments, respectively. **i,j**, The time evolution of bioluminscent flux signal evaluating the impact of total dose (**i**) and dose frequency (**j**). Schematics in (**a**), (**b**) were generated with BioRender. Data are presented as mean ± S.D. One-way ANOVA with Tukey’s post-hoc test for **c**,**d**,**g**,**h**. (* *P* ≤ 0.05, ** *P* ≤ 0.01, **** *P* ≤ 0.0001).

Local phage therapy significantly reduced bacterial burden compared to PBS controls, with mean log reduction in CFU/mg of 1.61, 2.33, and 2.37 for LD, HD, and DD, respectively **(Fig. 2d)**. Importantly, a substantial fraction of the mice in some of the conditions exhibited CFU readings below the limit of detection, indicating complete bacterial eradication. The daily dosing strategy achieved the highest eradication rate of PAO1-Xen41 (29%), surpassing LD (16%) and HD (18%) **(Fig. 2e).** These findings were supported by BLI **(Fig. 2f)**, which further revealed significantly lower flux in HD-treated mice compared to LD on Day 3 (**Fig. 2g**) and a trend toward reduced flux in DD-treated mice (**Fig. 2h**). Flux measurements over time indicated initial bacterial suppression followed by gradual regrowth in later days (**Fig. 2i, j**).

Together, these results demonstrate that localized free phage delivery, unlike IV or repeated IP delivery, effectively reduces bacterial burdens in wounds and, in some cases, achieves complete eradication of culturable bacteria in this model. The data further suggest that higher doses and repeat dosing might be more effective at reducing initial bacterial burden, which is in line with clinical practice.

### Development of an injectable HA-PEG hydrogel for sustained phage release

While our studies demonstrate the efficacy of localized free phage delivery using semi-occlusive dressings, this approach faces significant clinical limitations for two reasons. First, a semi-occlusive dressing cannot be used to retain a large volume of fluid in a human wound. Second, daily administration requires the patient to either be treated as an inpatient or to revisit the clinic daily, which is not reasonable for patients with limited mobility due to chronic wound infections. To address these limitations, we aimed to develop a hydrogel-based wound dressing incorporating phages with the potential to be used in existing clinical workflows, taking the once-weekly wound debridement appointments at the Stanford Advanced Wound Care Clinic as an example.

Hydrogels are well-suited as wound dressings. However, existing hydrogels typically deliver a low dose of phage (between 10^8^ to 10^10^ PFU/mL; lower than the LD treatment investigated earlier) in a burst-release (minutes to hours)^37–39^, we sought to develop a hydrogel that could deliver a high total dose of phage (10^11^ PFU/mL; on the order of the HD or DD treatment) in a sustained fashion over one week. Such a hydrogel would need to be compatible with both phage and the wound milieu, injectable, and capable of delaying the diffusive release of phage to deliver high titres over one week.

We developed a soft, injectable hyaluronic acid (HA)- and polyethylene glycol (PEG)- based hydrogel for PT. This hydrogel is synthesized through a crosslinking mechanism involving covalent thioether bonds between hyperbranched PEG multi-acrylate (HBPEG) and thiolated hyaluronic acid (HA-SH) as well as dynamic covalent hemithioacetal bonds between 4-arm PEG- aldehyde or PEG-benzaldehyde crosslinkers (4ALD and 4BLD) and HA-SH **(Fig. 3a).** These reactions occur spontaneously under physiological conditions, avoiding the need for harsh solvents or catalysts. Lytic phages can be mixed with the polymer precursors prior to gelation, and the resulting phage-hydrogel is injectable through a fine 29-gauge needle **(Fig. 3b)**.

**Fig. 3.**
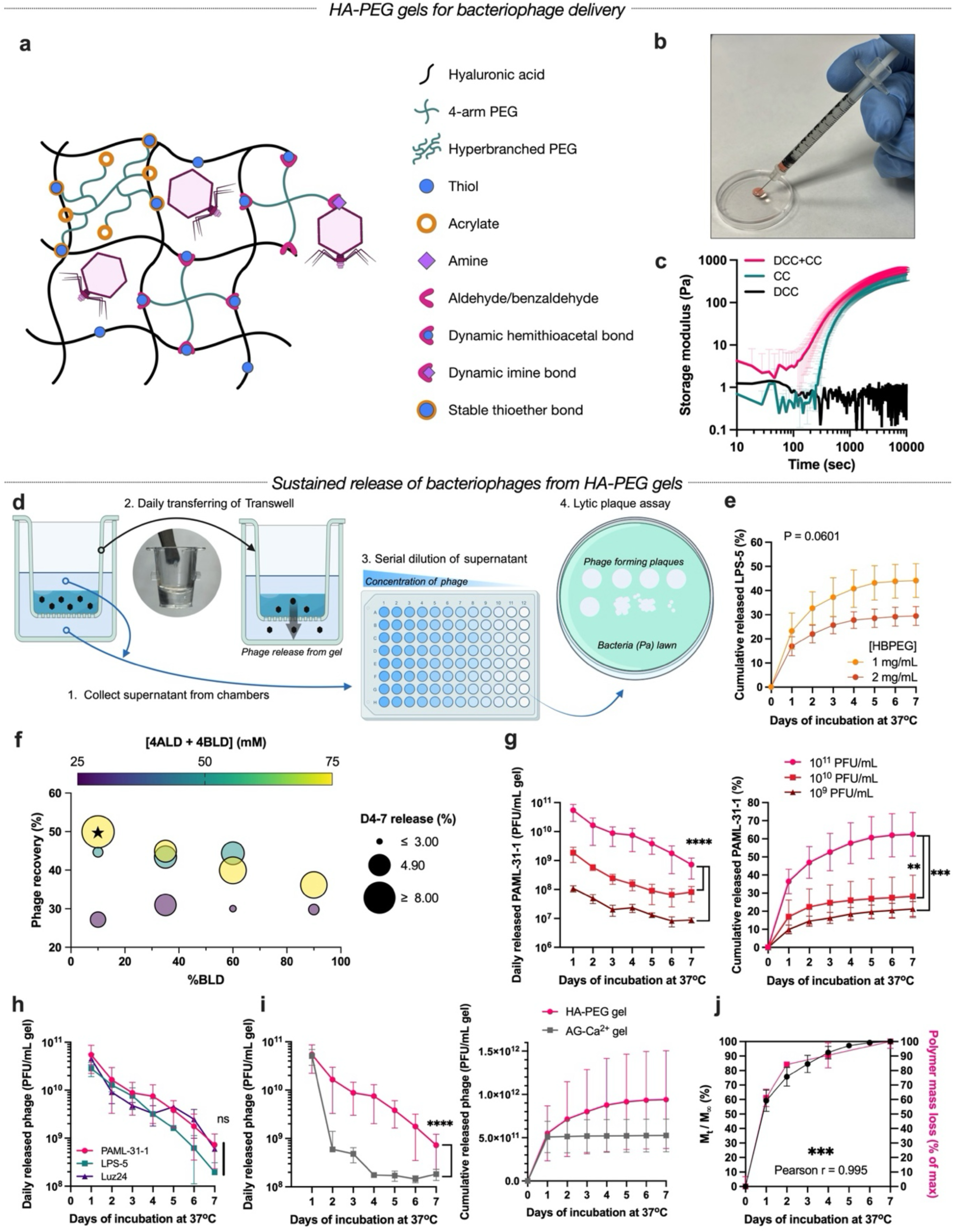
HA-PEG hydrogels for sustained delivery of high-dose bacteriophage. **a**, Schematic of hyaluronic acid (HA)- and polyethylene glycol (PEG)-based hydrogel (HA-PEG) crosslinking chemistries. Covalent thioether crosslinks are formed between thiolated-HA (HA-SH) and hyperbranched PEG multi-acrylate (HBPEG). Dynamic covalent hemithioacetal crosslinks are formed between HA-SH with either 4-arm PEG-aldehyde (4ALD) or benzaldehyde (4BLD). Amines presented on the surface of phages could theoretically bind to aldehyde/benzaldehyde through dynamic covalent imine bonds. **b**, Image of HA-PEG hydrogel injected through a 28G needle. **c**, The storage modulus of hydrogels with both dynamic covalent crosslinking and covalent crosslinking (DCC+CC, magenta), covalent crosslinking alone (CC, green), and dynamic covalent crosslinking alone (DCC, black). Hydrogel formulations: 7 mg/mL HA-SH, 4 mg/mL HBPEG, and 75 mg/mL of 4-arm PEGs with 10% 4BLD and 90% 4ALD for DCC+CC gel, n=4; 7 mg/mL HA-SH and 4 mg/mL HBPEG for CC gel, n=3; 7 mg/mL HA-SH and 75 mg/mL of 4-arm PEGs with 10% 4BLD and 90% 4ALD for DCC gel, n=1. **d**, Schematic of the transwell-release assay to quantify phage release from HA-PEG hydrogel daily. **e**, HBPEG of 1 mg/mL led to a higher percentage of phage recovery compared to 2 mg/mL; n=3 for each formulation. **f**, The HA-PEG hydrogel contained 5 mg/mL of HA-SH and 1 mg/mL of HBPEG, and the 4-arm PEGs concentration and %BLD were varied to explore the optimal hydrogel formulation with the highest percentage of recovery and sustained release. Sustained release was defined as the cumulative phage release from Day 4 to 7, indicated by the size of the dots; n=1 for each formulation with technical triplicates. The star sign indicates the chosen hydrogel formulation for the rest of the study. **g**. Quantification of daily and cumulative release of PAML-31-1 phage out of HA-PEG hydrogel with different encapsulation titres: 10^11^, 10^10^, and 10^9^ PFU/mL; n=8, 3, and 3, respectively, each with technical triplicates. **h**, Quantification of daily release of different phages— PAML-31-1, LPS-5, and Luz24—from HA-PEG hydrogel with encapsulation titre of 10^11^ PFU/mL; n=8, 4, and 2, respectively. **i**, Comparison of the daily and cumulative release of phage from HA-PEG hydrogel versus an ionically crosslinked alginate hydrogel (AG-Ca^2+^) with encapsulation titre of 10^11^ PFU/mL; n=8 and 4, respectively, each with technical triplicates. **j**, Close correspondence between the percentage of maximum phage release, defined as the ratio of cumulative phage release at given time point (M_t_) to the endpoint maximum cumulative release (M_∞_), and the mass loss of HA-PEG hydrogel, defined as the percentage of mass loss to the maximum mass loss; n=8 and 3, respectively. The PAML-31-1 phage release from HA-PEG gel data presented in (**h-j**) are referenced from the 10^11^ PFU/mL data in (**g**). Schematics in (**a**), (**d**) generated with BioRender. Data are presented as mean ± S.D for (**c**), (**e**), (**g**), (**h**), and (**j**). Repeated measures two-way ANOVA with Geisser-Greenhouse correction for (**e**), (**g**), (**h**) and (**i**) (PAML-31-1 vs LPS-5). Two-way ANOVA with mixed-effects model and Geisser-Greenhouse correction where phage type and day as fixed effects and gel ID as a random effect for (**i**) (PAML-31-1 vs Luz24 and LPS-5 vs Luz24). Bonferroni’s correction for multiple comparisons for (**g**) and (**h**). Pearson correlation for (**j**) (r = 0.995). (** *P* ≤ 0.01, *** *P* ≤ 0.001, **** *P* ≤ 0.0001).

The combination of static covalent crosslinking (CC) and dynamic covalent crosslinking (DCC) results in a higher and more rapid increase in stiffness compared to hydrogels with CC or DCC alone **(Fig. 3c)**. In addition, the fast kinetics of the DCC enables immediate enhancement of viscosity and adhesiveness without compromising injectability **(Extended Data Fig. 4a, b).** These features ensure precise application and secure adherence of the hydrogels to the irregular contours of infected wounds. This HA-PEG hydrogel’s unique properties make it highly suitable and easy to use for delivering therapeutic agents to wound beds.

We measured the release kinetics of phages from HA-PEG hydrogels *in vitro*, beginning with PAML-31-1. Hydrogels were formed in transwell inserts, immersed in a minimal amount of buffer, and the phage titre from the combined basolateral and apical chambers was measured using standard lytic plaque assays. The transwells were then transferred to a new well containing fresh buffer **(Fig. 3d)**.

To meet clinical needs, we optimized our hydrogel formulation for sustained phage release over one week. Prior work showed that hydrogels formed via thiol-Michael addition require polymer degradation mechanisms such as protease-cleavable crosslinkers to facilitate phage release^40^. We hypothesized that a lower thioether crosslinking density might enhance phage release by creating a looser polymer mesh, eliminating the need for additional polymer degradation mechanisms. Indeed, lower HBPEG concentrations increased phage recovery, suggesting that denser covalent thioether crosslinks trap phages and limit their release **(Fig. 3e and Extended Data Fig. 4c)**. Further investigations into hydrogels with low thioether crosslinking (1 mg/mL HBPEG) revealed that higher concentrations of 4ALD and 4BLD crosslinkers improved phage recovery **(Fig. 3f)**. At high concentrations of 4ALD and 4BLD, a lower percentage of benzaldehyde (%BLD) crosslinkers corresponded to higher percent phage recovery and more sustained release from day 4 to 7, likely due to more stable polymer-polymer and polymer-phage interactions with benzaldehyde compared to aldehyde. Thus, we identified that the optimal HA-PEG hydrogel formulation for sustained phage release was 5 mg/mL HA-SH, 1 mg/mL HBPEG, and 75 mg/mL of 4-arm PEGs with 10% 4BLD and 90% 4ALD, which was used for the remainder of the study and is referred to as HydroPhage (HP).

Next, we assessed how phage titre and identity affected release kinetics. Higher encapsulated phage titres increased recovery; encapsulating 10^11^ PFU/mL in HP resulted in 62% phage recovery, compared to 28% recovery for 10^10^ PFU/mL and 21% recovery for 10^9^ PFU/mL **(Fig. 3g)**. Higher titres may saturate the binding of aldehyde and benzaldehyde groups, allowing a larger portion of unbound phages to diffuse freely. Testing three different *P. aeruginosa*-targeting phages (Luz24, PAML-31-1, and LPS-5) revealed comparable release kinetics **(Fig. 3h)**. Unlike an ionically crosslinked alginate (AG-Ca^2+^) hydrogel, which exhibited a burst release of nearly all encapsulated phages within the first 24 hours, HP exhibited sustained high-titre phage release (∼10^9^ PFU/mL or higher) for up to 7 days **(Fig. 3i)**.

Finally, we explored the mechanisms driving phage release. While dynamic covalent imine bonds between phage surface amines and aldehyde/benzaldehyde groups could theoretically sustain phage release from HP^41–43^, and benzaldehyde groups are expected to form bonds with higher affinity than aldehyde groups^62,63^, varying the ratio of aldehyde to benzaldehyde had little effect on phage release. Instead, phage release closely correlated with polymer mass loss **(Fig. 3j)**, indicating hydrolytic erosion as the primary mechanism. A fluorescent bead diffusion assay estimated the hydrogel mesh size between 20-200 nm **(Extended Data Fig. 4d)**, aligning with the 174 nm length and 61 nm head diameter of PAML-31-1 phage **(Fig. 1b)**. At 18 hrs, 30-35% of phages were released **(Extended Data Fig. 4e)**, further confirming this estimate. Overall, our dual-crosslinked HA-PEG hydrogel offers a novel approach for sustained high-titre phage delivery.

### HP disrupts *P. aeruginosa* biofilms

We conducted *in vitro* tests to assess the antibacterial properties of HP. For HP encapsulating 10^11^ PFU/mL, the amount of PAML-31-1 released on Day 7 suppressed 30% of planktonic *P. aeruginosa*, highlighting the extended antibacterial effect **(Fig. 4a)**. This prolonged suppression underscores the dose-dependent nature of phage-bacterial interactions and the potential of HP to control bacterial burden over extended timeframes *in vivo*.

**Fig. 4.**
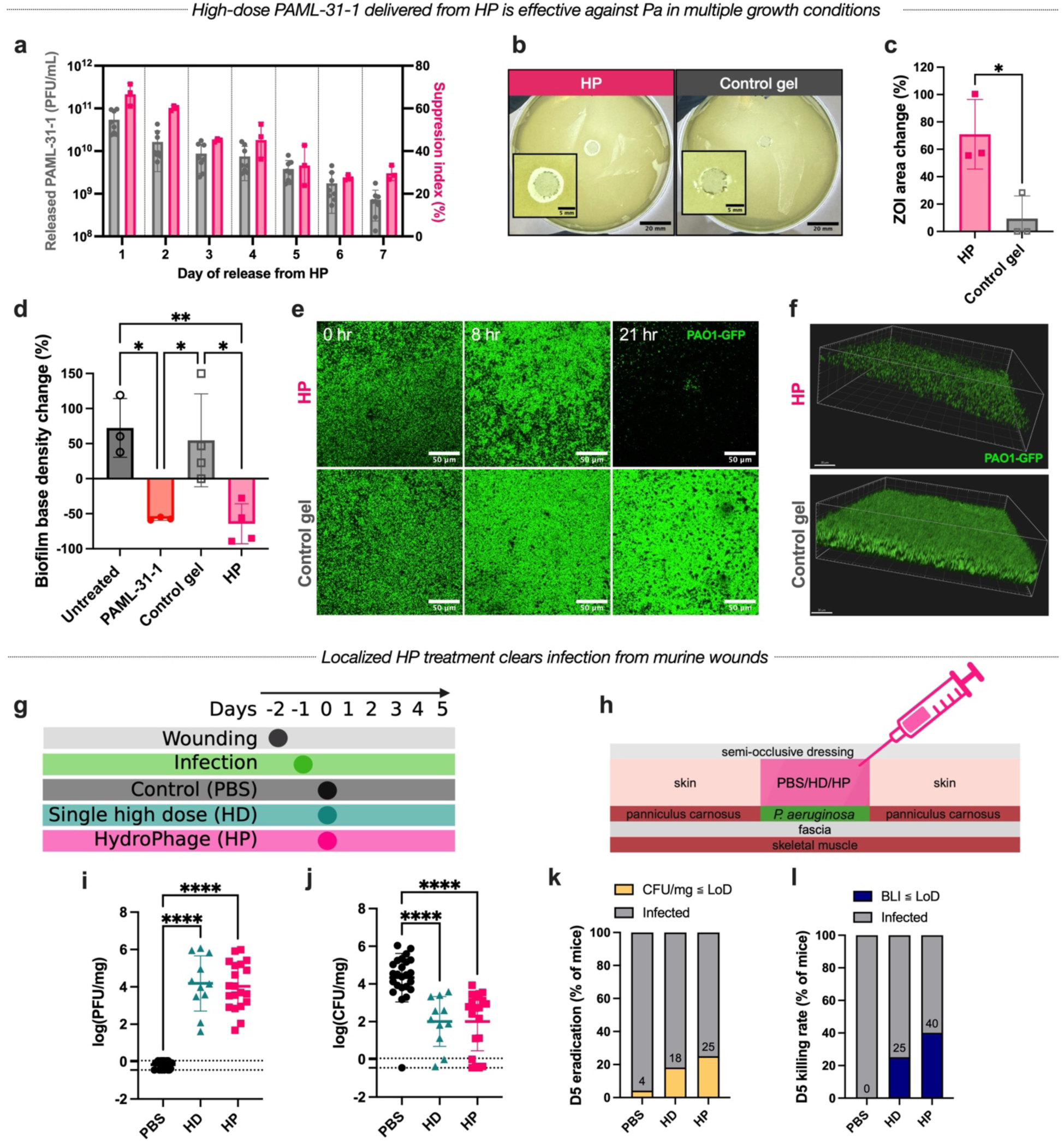
Phage-delivering HA-PEG hydrogels (HydroPhage, HP) suppress *P. aeruginosa in vitro* and clears wound infections *in vivo*. a, Quantification of suppression index (SI%) against PAO1-Xen41 by planktonic suppression assay (PSA) (n=3, pink) with the PAML-31-1 titre released from HP in transwell-release assay daily (n=8, gray). The daily released PAML-31-1 titre data are referenced from Figure 3g. **b**,**c**, The zone of inhibition (ZOI) formed by HP and control gel at 24 hours of gel placement on *P. aeruginosa* lawn. The representative images of HP and control gel on *P. aeruginosa* lawn (**b**), and quantification of change in ZOI area at 24 hours as a percent of initial area of hydrogel (**c**); n=3 gels and n=2 independent experiments each. **d**,**e**,**f**, PAO1-GFP biofilm was formed on the surface of top agar and treated with untreated control, PAML-31-1 phage in media, control gel or HP for 21 hours. The quantification of biofilm base density changes (**d**), representative confocal images of biofilm incubated with HP and control gel treatments over hours (**e**), and the 3D rendering of biofilm at 21 hours (**f**); n=3, 3, 4, and 4 replicates for untreated, PAML-31-1, control gel, and HP groups respectively; n = 2 independent experiments each. **g**, Illustration of a murine wound infection study to test the efficacy of HP and IV phage injection treatments. **h**, Illustration of how mice receive HP or PBS control treatment by topical injection under Tegaderm dressing. **i**,**j**,**k,l**, Endpoint characterization of harvested mice wound tissue. The phage counts (PFU/mg) (**i**) and bacterial burden (**j**) receovered from mouse tissue. n=24, 11, 20 mice and n=5, 2, 3 independent experiments for PBS, HD, and HP groups, respectively; The eradication rate for PBS, HD, and HP groups (**k**). n=24, 11, 20 mice and n=5, 2, 3 independent experiments, respectively; the killing rate for PBS, HD, and HP groups (**l**). n=17, 12, 15 mice and n=4, 2, 3 independent experimetns, respectively. Data are presented as mean ± S.D. Unpaired two-tailed Student’s t-test for (**c**). One-way ANOVA with Tukey’s post-hoc test for (**d**), (**i**), and (**j**). Scale bars: 20 mm for (**b**) and 5 mm for the inset, 50 µm for (**e**), and 30 µm for (**f**). (* *P* ≤ 0.05, ** *P* ≤ 0.01, **** *P* ≤ 0.0001).

Next, we evaluated if PAML-31-1 could escape HP and lyse *P. aeruginosa* on a semi-solid substrate without fluid. On Mueller-Hinton agar plates inoculated with *P. aeruginosa*, HP increased the zone of inhibition (ZOI) by 60% compared to a control gel, showing that phages could escape HP under drier conditions and propagate beyond the area in direct contact with HP **(Fig. 4b, c)**.

We also tested HP’s efficacy against biofilms, which are dense polymeric matrices that harbor active and dormant bacterial populations, are prevalent in chronic wounds^44,45^, and diminish the efficacy of antimicrobials, complicating treatment^46–50^. Biofilms of PAO1-GFP were grown on top agar and treated with PAML-31-1 in media, HP, or a control gel in a transwell insert after 24 hours. Both PAML-31-1 and HP reduced biofilm thickness by 43% and 52%, respectively, at 21 hours post-treatment, with no disruption of biofilm growth observed with the control gel or untreated samples **(Extended Data Fig. 6a)**. Biofilm density also decreased by 57% with PAML-31-1 and 65% with HP, while the control gel and untreated control exhibited increases in biofilm density of 55% and 72%, respectively **(Fig. 4d)**. Confocal microscopy vividly illustrated marked to complete disruption under treatment with PAML-31-1 and HP **(Fig. 4e, f and Extended Data Fig. 6b)**. In addition, PAML-31-1 in media and HP also significantly reduced biofilm density of an extensively drug resistant (XDR) clinical *P. aeruginosa* isolate we collected from Stanford Healthcare Clinical Microbiology Laboratory (SHCML) (**Extended Data Fig. 5c, d)**^51^. Together, these results demonstrate that HP effectively suppresses and disrupts *P*a under both planktonic and biofilm conditions.

### Localized HP reduces the bacterial burden in infected murine wounds

To evaluate the *in vivo* efficacy of the HP, we compared its performance to PBS and high-dose (HD) free phage treatments **(Fig. 4g)**. Mice received HP as a topical treatment or HD topical free phage, with the same total amount of phage administered **(Fig. 4h)**. Similar amounts of phage were recovered in the wound homogenate between HD and HP **(Fig. 4i)**. Mice treated with HP demonstrated a significant >2-log reduction in CFU/mg wound tissue relative to PBS (2.33, n=20), comparable to the effect of HD (2.33, n=11) **(Fig. 4j)**. Treatment with HP resulted in an eradication rate of 25%, compared to 18% for HD **(Fig. 4k)**. BLI results corroborated these findings **(Fig. 4l)**. These results demonstrate that a single injection of HP can effectively reduce the bacterial burdens in mice wounds and eradicate infections as well as or better than a single high-dose of free phage, highlighting the merit of local, sustained delivery of high-titre phages via hydrogels.

### Impact of successful treatment with HP on cellular inflammation

Pathogenic bacteria, especially *P. aeruginosa*, drives persistent cellular inflammation in chronic wounds. To determine whether successful treatment with HydroPhage (HP)—defined as cases where HP eradicated *P. aeruginosa*—was associated with changes in cellular inflammation, we characterized the cutaneous immune response to *P. aeruginosa* infection and its modulation by HP treatment.

Using a 13-color flow cytometry panel **(Extended Data Table 1)**, we quantified the abundance of CD45^+^ cells, natural killer (NK) cells, neutrophils, macrophages, B cells, and T cells in infected wounds, while excluding monocytes, monocyte-derived dendritic cells (MoDCs), and conventional dendritic cells (cDCs) based on the gating strategy shown in **Extended Data Fig. 6**. Subpopulations of innate lymphocytes were not further discriminated.

*P. aeruginosa* infection significantly increased CD45^+^ cells and NK cells **(Fig. 5a, b)**. Neutrophils were found to be increased in infection while macrophage and T cells and were found to be reduced **(Fig. 5c-e).** Reduced proportions of CD45^+^ cells, NK cells, and neutrophils following successful eradication of *P. aeruginosa* by HP treatments, possibly suggesting reduced cellular inflammation. Macrophage and T cell abundance remained low even in cases where bacteria were eradicated **(Fig. 5e, f)**.

**Fig. 5.**
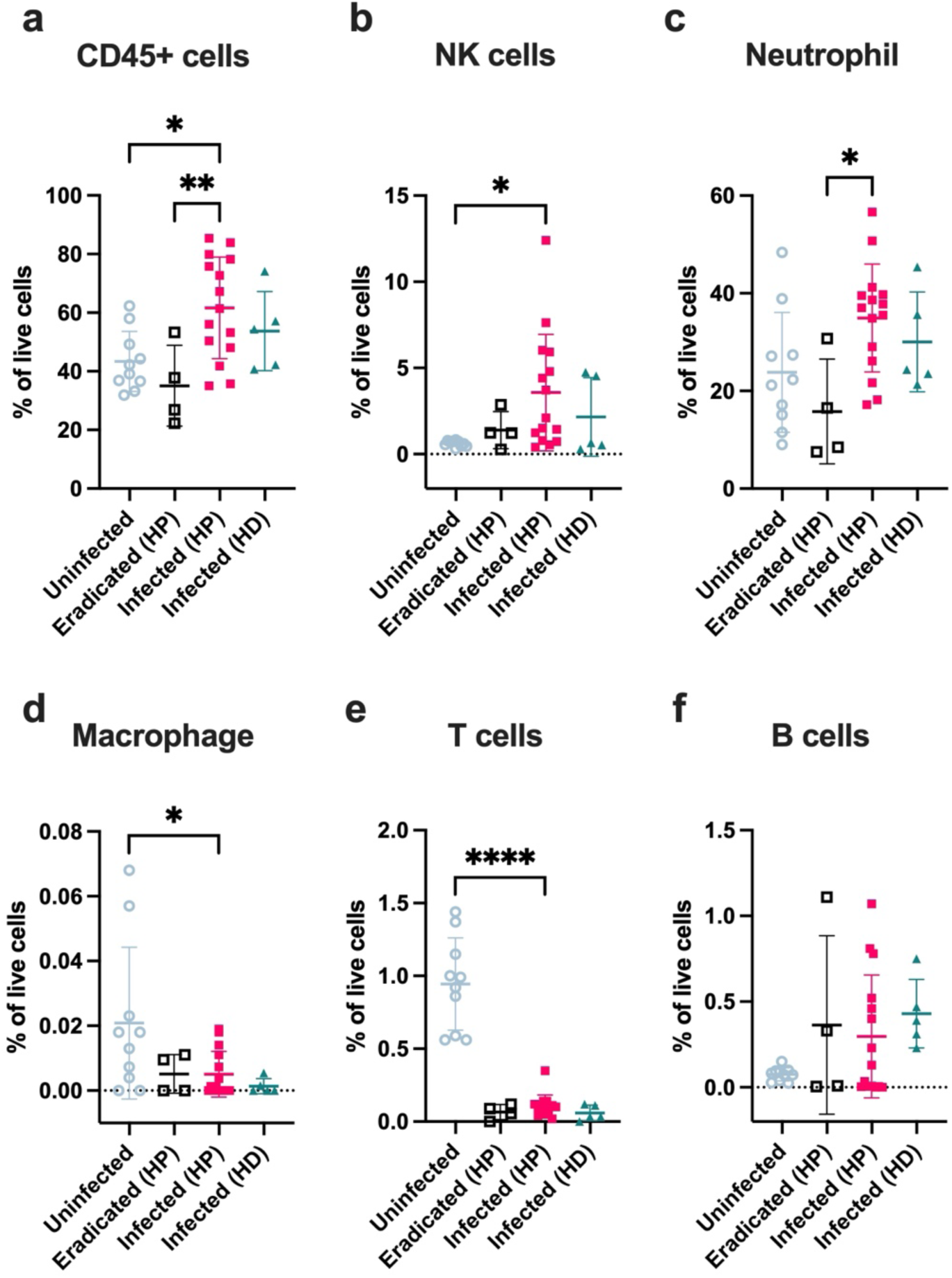
HydroPhage treatment reduces skin-associated immune cell populations. **a-f**,The abundance of CD45^+^ (**a**), natural killer (NK) cells (**b**), neutrophils (**c**), macrophages (**d**), T cells (**e**), and B cells (**f**) in uninfected mouse wounds versus wounds infected with *P. aeruginosa* and treated with HP successfully (Eradicated (HP)), or which remained infected (Infected (HP)), or which remained infected after treatment with HD (Infected (HD)). n=10, 4, 15, 5 mice and n=1, 3, 3, 2 independent experiments for each group, respectively. See **Extended Data** Figure 6 for gating strategy and cell definitions. Data are mean ± S.D. Ordinary one-way ANOVA with Tukey post-hoc tests for (**a-f**) and with Dunnett post-hoc tests versus Infected (HP). (* *P* ≤ 0.05, *** *P* ≤ 0.001, **** *P* ≤ 0.0001)

We also evaluated the biocompatibility and immunogenicity of the HA-PEG hydrogel and towards RAW264.7 murine macrophage-like cells grown *in vitro* **(Extended Data Fig. 7a)**. Neither the control gel nor PAML-31-1 exhibited any cytotoxicity **(Extended Data Fig. 7b)** or induced M1/M2 polarization **(Extended Data Fig. 7c)**. While limited in scope, these *in vitro* results suggest that the HA-PEG hydrogel is biocompatible and non-immunogenic towards macrophages.

### Resistance to PAML-31-1 develops independently of treatment dosage, route, and efficacy

Despite the success of HP and DD in eradicating infections, eradication rates did not exceed 30% **(Fig. 6a)**. We hypothesized that bacteria might develop resistance against phages if not fully eradicated, similar to behavior seen *in vitro* **(Extended Data Fig. 1a)**. The positive correlation between phage and bacteria counts at the endpoint suggested that persistent bacterial burdens were not due to phage depletion **(Fig. 6b, c)**.

**Fig. 6.**
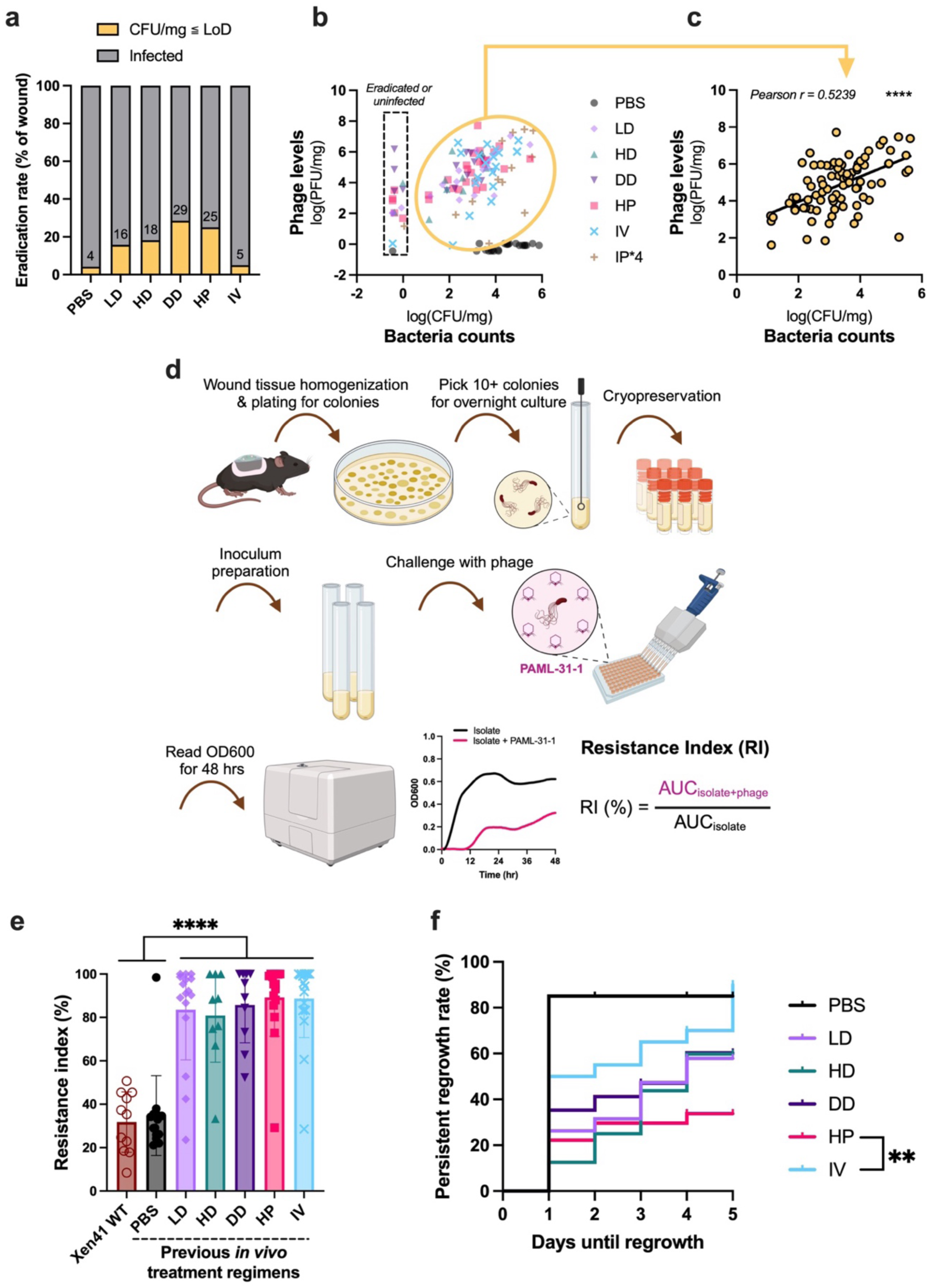
Development of phage resistance from PT is independent of treatment route, dosing, and efficacy. **a,** Summary of eradication rates by phage treatments investigated in this study. The isolates from endpoint infected wound were used for downstream resistance characterization. **b,** The endpoint phage counts recovered from mice wounds were plotted against the bacteria counts recovered. The dotted box encircles the wounds that did not have bacteria counts, either from eradication or not infected. **c**, The correlation of log-transformed PFU/mg to CFU/mg for the wounds that had both phage and bacteria recovered; n=88. **d,** Schematic of PSA for the mice wound isolates to characterize bacteria resistance after prior PAML-31-1 treatment regimens, compared to PBS-treated control and wild-type PAO1-Xen41. **e,** Quantification of resistance index (RI%) for PAML-31-1 against PAO1-Xen41 WT and wound isolates with previous *in vivo* treatment regimens; n=11 for PAO1-Xen41 WT, n=15 for PBS, n=19 for IV, n=16 for v, n=9 for HD, n=10 for DD, and n=16 for HP. **f**, The persistent regrowth rate of different treatment regimens, HP is the only localized treatment presents significant difference with IV treatment (P = 0.0064). Dots on the curves indicate mice were censored due to Tegaderm detachment or were still uninfected on Day 5. Schematic in (**c**) was generated with BioRender. Data are mean ± S.D. Two-tailed Pearson r correlation for (**b**). One-way ANOVA with Tukey’s post-hoc test for (**e**). Log-rank test with Bonferroni’s correction for multiple comparison for (**f**). (** *P* ≤ 0.01, **** *P* ≤ 0.0001).

To confirm the development of resistance to PAML-31-1 *in vivo* in response to phage treatments, we re-challenged stationary phase bacterial isolates from previous *in vivo* treatments with PAML-31-1 **(Fig. 6d)**. Bacterial isolates from all phage-treated groups showed significant increase in resistance index (RI) upon re-challenge compared to phage-unexposed wild-type PAO1-Xen41 and PBS control groups, indicating that substantial regrowth of resistant bacteria had occurred, even with ineffective treatments such as IV injection **(Fig. 6e)**. The comparable RI% among resistant isolates across treatment groups suggests that the degree of resistance was independent of treatment efficacy.

*Pseudomonas* isolates from HydroPhage-treated mice that showed complete resistance in the PSA and EOP **(Extended Data Fig. 8a, b)** were selected for whole genome sequencing. In these cases, we identified nonsynonymous mutations predicted to encode missense variants in LPS biosynthesis proteins. As OSA/LPS is the PAML-31-1 receptor, this data suggests that treatment with HP selects for PAML-31-1-resistant mutants **(Extended Data Fig. 8c)**.

We next assessed which treatments were most effective at preventing bacterial regrowth over time. For this, we compared the percentage of mice with wounds having persistent bacterial regrowth, as assesssed from BLI measurements **(Fig. 1h)**. Interestingly, among localized treatments, only HP exhibited significantly suppressed bacterial regrowth compared to IV treatment **(Fig. 6f)**.

These results show that PAML-31-1 resistance develops regardless of treatment dosage, frequency, delivery route, and was not correlated with the reduction of bacterial burden.

## Discussion

In this study, we investigated the effectiveness of systemic and topical PT, including a custom-designed hydrogel capable of sustained phage release, using a mouse model of stable *P. aeruginosa* wound infection that we adapted for study of PT.

Our findings demonstrate rigorously that local, topical delivery is superior to systemic (single IV or repeated IP) delivery, confirming a principle that is anecdotally appreciated but lacks experimental support. It remains possible that systemic PT may have a role in reducing the systemic dissemination of bacteria or progression to sepsis. However, such an effect was not explored in this model that focuses on evaluating bacterial burden reduction and emergence of resistance in wounds. Despite repeated localized dosing (DD) resulting in similar bacterial burden at the end of our experiment compared to single low-dose (LD), we observed greater suppression of initial burden through real-time BLI and higher eradication rates at the study endpoint. Further investigation is needed to determine if higher doses or repeated dosing are more effective than lower doses at suppressing bacterial burden beyond our monitored period. Nonetheless, our findings provide experimental support for current anecdotal guidelines from the Antibacterial Resistance Leadership Group (ARLG) Phage Taskforce in the USA that recommend high doses, repeated administration, and local PT when feasible^52^.

Building on these insights, and recognizing that using a semi-occlusive dressing to trap free phage at the wound infection is clinically unfavorable in most cases, we developed “HydroPhage”, a hydrogel system that delivers high-titre phages in a sustained fashion over one week. Eradication of infection was associated with decreased CD45^+^ cells, NK cells, and neutrophils, suggesting that successful treatment with HP can decrease certain types of cellular inflammation seen in infected wounds, although further investigation is warranted. Our results offer a new phage delivery platform that is biodegradable, adaptable to wound treatment, and highlights the significant potential of hydrogels in containing bacterial growth in wounds.

A key innovation was the use of hyaluronan-based DCC hydrogels for phage delivery, which enables the sustained release of phage at high titres (i.e., >10^9^ PFU/mL gel) over a week. Hydrogels are promising as wound dressings and platforms for controlled topical phage delivery^53–55^. Previous hydrogel- or microparticle-based approaches for phage delivery often present either rapid burst release^38,39,41,56–58^ or are clinically impractical due to low recovery of phage without mechanical (e.g., shaking) or biochemical (e.g., protease activity) interventions^40,59,60^. DCC hydrogels, with their intrinsic viscoelasticity, injectability, and self-healing characteristics, have gained significant attention in many biomedical applications^41,61–63^. Among various DCC chemistries considered, hemithioacetal crosslinking is particularly suited for topical applications, as its rapid kinetics and high reversibility provide immediate enhancement of viscosity and shear-thinning behavior^64^. We leveraged these properties, combined with the stability of covalent thioether bonds, to develop an adherent, slow-gelling HA-PEG hydrogel. The low covalent thioether crosslinking density provides stability without trapping sub-micron scale particles like phages, and allows for slow erosion, enabling sustained release of phages from the hydrogel. Additionally, HP is easily injectable and can conform to the irregular contour of the wound bed, ensuring prolonged contact with the affected area. Both HP and PT were safe in the mouse infection model, and successful bacterial eradication reduced certain immune cell populations, further supporting the safety of both HP and PT^55,65–67^. We propose that HP offers an effective and clinically practical way to deliver phages for treating MDR wound infections. We envision that PT should integrate into chronic wound care regimens in the form of an advanced dressing that delivers effective concentrations of phages over a sustained period.

Surprisingly, despite the clear advantage of topical over systemic PT in our model, we found that phage PFU levels within the wound bed were equivalent between systemic and topical treatments. This suggests that phage propagation within tissues may be insufficient for bacterial clearance and that effective phage therapy may require sufficiently high doses of phage to be delivered locally over a limited time window. This suggests that the pharmacokinetics of phage delivery play a critical role in treatment efficacy.

Our findings also highlight the challenge of phage resistance in PT and the potential limitations of phage monotherapy^68,69^. *In vitro* and *in vivo* studies demonstrated bacterial regrowth to single PT within 72 hours, with eradication achieved in only a minority of cases, irrespective of treatment strategies employed. Resistance was likely attributed to nonsynonymous mutations encoding missense variants in phage receptors. While phage cocktail treatments^70^ may mitigate this resistance, such approaches entail a level of complexity that was out of the scope of this investigation. Furthermore, there is a lack of comprehensive data supporting the superiority of cocktail therapy over monophage therapy, either in preclinical models or clinical trials.

The limitations of this study include the reliance on a single strain of *P. aeruginosa* (PAO1-Xen41) and a single phage (PAML-31-1) for the *in vivo* infection model. It would be important to extend these findings to include other pathogens and phages, including MDR *P. aeruginosa* clinical isolates, polymicrobial infections, and phage cocktails. The use of Tegaderm in this model was essential for delivering topical liquids (e.g., phages) and for preventing secondary infection of exposed wounds by environmental pathogens. However, by containing PT within the wound bed, this dressing may have blunted the potential advantage of repeated dosing or hydrogel-enabled sustained release. Similarly, the presence of adherent bacteria may have surreptitiously increased bacterial counts as assessed by luminescence. Nonetheless, the assays and models developed here can facilitate further investigations of PT. As configured, this model provides unique insights into resistance dynamics throughout the treatment course with paired endpoint analyses.

We conclude that these studies experimentally establish key treatment principles for PT in a mouse model. They provide a path forward for effective PT by enabling local, sustained, high-dose phage delivery using hydrogels.

## Methods

### Phage selection and propagation

Phages were propagated using methods previously established in the lab^71^. PAML-31-1, LPS-5, Luz24, and OMKO1 were propagated in *Pseudomonas aeruginosa* (*P. aeruginosa*) strain PAO1. Phages were chosen because they could be reliably propagated to high titers (>10^12^ PFU/mL) in PAO1. Early- to mid-log phase cultures were infected with phages to generate lysates from 300 mL cultures.

### Phage purification using AKTA-FPLC system

Bacteria were removed by 3 rounds of centrifugation at 8000 x *g*, 10 min, 4°C. Supernatants were filtered through a 0.22 μm polyethersulfone (PES) membrane filter (Corning, Cat. No. 431118). The supernatant was treated with 50 U/mL Benzonase Nuclease (Sigma-Aldrich, Cat. No. E8263) overnight to digest free DNA. Benzonase-treated lysates were then mixed with 3M KH_2_PO_4_ buffer (pH 7.0) in a 1:1 ratio in preparation for purification by fast protein liquid chromatography (FPLC). The FPLC-based virus purification was conducted with a CIMmultus OH 1 mL monolithic column with a 2 μm channel size (Sartorius BIA Separations, Ajdovščina, Slovenia). The washing, regeneration, performance test, and storage of the column was carried out according to manufacturer recommendations. The column was attached to an ÄKTA Pure FPLC (GE Healthcare Biosciences, Sweden) equipped with a P960 sample pump and analyzed with UNICORN 5.0 software (Cytiva Life Sciences, USA). All purification protocols were run at room temperature and all buffers and samples were filtered through a 0.22 µm PES membrane. 40-60 mL of the mixed sample was loaded onto a pre-equilibrated column with 1.5 M KH_2_PO_4_ loading buffer, pH 7.0. After sample loading, the column was washed with 10 column volumes of loading buffer to remove unbound particles. The elution of bacteriophage was performed using a linear gradient over 20 column volumes from 1.5 M to 20 mM KH_2_PO_4_. The fraction corresponding to the eluted phage was collected based on UV A280 measurement. We then dilute or perform buffer exchange on the eluted phages into Tris+Mg buffer (100 mM NaCl, 50 mM Tris-HCl, 8 mM MgSO_4_, pH 7.50).

### Phage characterization

The size and morphology of phages were assessed as was done previously using transmission electron microscopy (TEM) on a JEOL JEM1400 (JEOL USA Inc., Peabody, MA, USA) operating at 80 kV^71^. In brief, 5 µL of diluted phage solution was applied onto carbon-coated copper grids (FCF-200-Cu, Electron Microscopy Sciences, Hatfield, PA, USA). Following a 3-minute incubation, the grid was immersed in ddH2O and subsequently treated with 1% uranyl acetate for staining. After drying for 15 min, the sample was ready for TEM imaging.

### Hydrogel preparation

For making HA-PEG hydrogel, thiolated hyaluronic acid (HA-SH, 450 kDa, #HASH0102, Blafar Ltd, Ireland), hyperbranched PEG multi-acrylate (HBPEG, 10 kDa, #HBPEG10K, Blafar Ltd, Ireland), 4-arm PEG-aldehyde (4ALD, 10 kDa, #PSB-4303, CreativePEGWorks, USA), and 4-arm PEG-benzaldehyde (4BLD, 10 kDa, #6020710909, SINOPEG, China) were reconstituted in PBS+Mg (Phosphate-Buffered Saline + 8 mM MgSO_4_, pH 7.50) at concentration of 1wt%, 5wt%, 40 mM, and 40 mM respectively, adjusted to neutral pH, purged with argon gas and kept on ice before use. Appropriate volumes of the following reagents were added into an Eppendorf tube in the order of: PBS+Mg, HBPEG, 4BLD, 4ALD to make a crosslinker mixture. Next, phage was added to the crosslinkers and vortexed, then HA-SH was added, vortexed again, and mixed thoroughly by pipetting. The viscous slow-gelling mixture then was withdrawn by pipette to deposit into transwell inserts (Corning, USA, Cat. No. 353492) for *in vitro* assays or by insulin syringe with a 28G needle (BD, REF. 329461) to inject for *in vivo* treatment. The final formulation of HA-PEG hydrogel was 5 mg/mL HA-SH, 1 mg/mL HBPEG, 75 mg/mL of 4-arm PEGs with 10% 4BLD and 90% 4ALD, and 10^11^ PFU/mL phage unless otherwise noted.

For making the ionically crosslinked alginate hydrogel (AG-Ca^2+^), lyophilized alginate (PRONOVA UP VLVG, #4200501, IFF, USA) was reconstituted in Tris+Mg buffer (50 mM Tris-HCl, 100 mM NaCl, 8 mM MgSO_4_) at 4wt%. Calcium sulfate was prepared as a form of saturated slurry in deionized water at 500 mM. Reconstituted alginate and phage solution were added to a 1 mL syringe, and the calcium slurry was added into another syringe using a pipette. The solution in each syringe was thoroughly mixed by moving the plunger up and down before the syringes were connected with a Luer Lock. Alginate, calcium, and phages were then mixed thoroughly between syringes and 100 μL was deposited into a transwell insert. The formulation of the AG-Ca^2+^ hydrogel was 2wt% alginate, 50 mM calcium, and 10^11^ PFU/mL phage.

### Rheological measurements

Rheometry was performed on Discovery HR-2 hybrid rheometer (TA Instruments). The rheometer was heated to 37°C before measurements. 60 µL of hydrogels were deposited directly onto the rheometer base plate. An 8-mm flat plate was then immediately lowered to contact the gel and form an 8-mm disk. Mineral oil (Sigma) was applied to seal the edges of the gel to prevent dehydration. Time sweep measurements were performed at 1 rad s^-1^ and 1% strain until the storage and loss moduli plateaued.

### Transwell-release assay

Transwell inserts (Falcon, Corning, USA, Cat. No. 353492) for 24-well plates were blocked in a solution of 2% bovine serum albumin (BSA) in PBS for 1 hour, rinsed with PBS+Mg for 10 min, thoroughly aspirated, and then air-dried for 20 min at room temperature prior to use. 100 µL of hydrogel with encapsulated phage was aliquoted into each transwell insert and incubated in a closed box at room temperature for 3.5 hours until the hydrogels had fully gelled. After gelation, transwells were transferred to 24-well tissue culture-treated plates (Corning, USA, Cat. No. 3524) containing 500 µL of PBS+Mg buffer. An additional 200 μL of PBS+Mg was added on top of the hydrogel to prevent drying. After 24 hours of incubation at 37°C, the transwell insert with hydrogel was removed. The buffer on top of the hydrogel and any residual buffer on the bottom of the transwell were carefully collected and combined with the buffer in the 24-well plate and stored at 4°C for phage titre quantification. The transwell was transferred into a new well containing 500 μL of fresh buffer, and 200 μL of fresh buffer was added on top. This process was repeated for 7 days.

### HA-PEG hydrogel erosion

The erosion study on HA-PEG hydrogel without phage encapsulation was conducted by forming hydrogels in the transwell insert, submerging in PBS+Mg buffer, and transferring daily as described above. The dry mass of the hydrogel was measured at each time point after lyophilization. Erosion, defined as the percentage to maximum polymer mass loss, was quantified as the loss of dry mass at each time point normalized to the endpoint of the assay on Day 7.

### Beads diffusion assay

Fluorescent beads of 20 nm (FluoSpheres carboxylate, ThermoFisher Scientific, Cat. No. F8783) and 200 nm (FluoSpheres carboxylate, ThermoFisher Scientific, Cat. No. F8807) were encapsulated in HA-PEG hydrogels (5 mg/mL HA-SH, 1 mg/mL HBPEG, 75 mg/mL of 4-arm PEGs with 10% 4BLD and 90% 4ALD) at a concentration of 2 mg/mL during the 3.5-hour gelation period in the Transwell inserts (Falcon, Corning, Cat. No. 353492). The hydrogels in Transwell were transferred to fresh wells containing 500 µL of PBS+Mg buffer in a 24-well tissue culture-treated plates (Corning, USA, Cat. No. 3524), and additional 200 µL were added on the top. The plates were then incubated at 37°C in a sealed container. At 3 and 18 hours, the supernatants from the top and bottom compartments were collected, combined, and 50 µL of each sample was transferred to a 96-well flat-bottom plate (Falcon, Corning, USA, Cat. No. 351172) for fluorescence measurement using a microplate reader (Spark, Tecan, Swiss). The percentage of released beads was calculated from the fluorescence intensity of the supernatant, normalized to the initial encapsulation and referenced against standard curves generated for each bead size.

### Plaque assays

Phages were enumerated by plaque assays using the spot dilution double-agar overlay technique. 100 μL of mid-log phase bacteria was added to 5 mL of top agar (5 g/L agar, 10 g/L tryptone, 10 g/L NaCl, 20 mM MgSO_4_, 20 mM CaCl_2_) and poured onto LB-agar plates to solidify. 10-fold serial dilutions of phages were prepared in 96-well polystyrene U-bottom tissue culture-treated plates (ThermoFisher Scientific, USA, Cat. No. 168136) in SM buffer (50 mM Tris-HCl, 100 mM NaCl, 8 mM MgSO_4_, 0.01% gelatin (Sigma-Aldrich, USA, Cat. No. G7041), pH 7.50). 10 μL of each dilution was spotted onto the top agar, incubated at 37°C overnight, and plaques were counted.

### Planktonic suppression assay (PSA)

Overnight cultures were prepared from a glycerol stock of PAO1-Xen41 or from cryopreserved mice wound isolates by directly inoculating LB medium and incubating at 37°C with shaking at 200 x *rpm* overnight. A 50 μL aliquot of diluted overnight culture in “modified-LB” (LB buffer + 50 mM Tris-HCl + 8 mM MgSO_4_, pH 7.50) was prepared at the concentration of 3.75 x 10^7^ CFU/mL in a 96-well flat-bottom plate (Falcon, Corning, USA, Cat. No. 351172). 100 μL of phage suspension in modified-LB was added to each well containing bacteria to reach the target MOI in a total volume of 150 μL. Wells were then topped with 15 μL of mineral oil and covered with a gas-permeable sealing membrane to prevent evaporation (USA Scientific, Cat. No. 9123-6100). The final co-cultures were incubated at 37°C in an automated spectrophotometer (Biotek Synergy 2, Agilent Technologies, USA) for 48 hours, with OD600 measurements taken every 20 min following 5 seconds of shaking. The control consisted of bacteria incubated with modified-LB without phages. Each sample was prepared in triplicate. The Suppression Index (SI%) was calculated using the formula provided below to quantify the relative suppression of bacterial growth in phage-treated versus non-treated condition^35^.

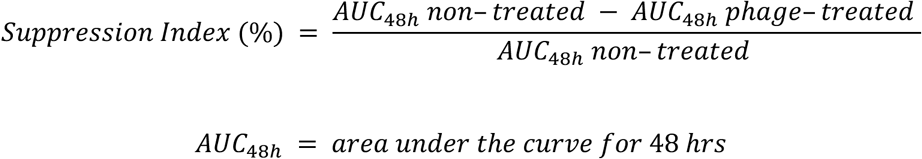

Phage titres were measured using plaque assays against *P. aeruginosa* PAO1. To evaluate the efficiency of different phages to suppress planktonic PAO1-Xen41, an MOI of 8,000 was compared. To evaluate the resistance of mouse isolates to PAML-31-1 phage, resistance index was tested at an MOI of 500.

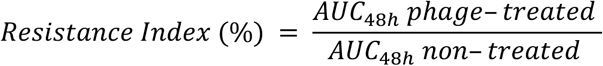

### Zone of inhibition (ZOI) assay

An overnight culture of PAO1 was subcultured to an OD600 of 0.65, then 100 µL was spread onto a Mueller-Hinton agar plate and air-dried. 100 µL of pre-formed HA-PEG hydrogel (5 mg/mL HA-SH, 1 mg/mL HBPEG, 75 mg/mL of 4-arm PEGs with 10% 4BLD and 90% 4ALD, and 10^11^ PFU/mL PAML-31-1 phage) was transferred onto the inoculated agar plate after 3.5 hours of gelation. The hydrogel was allowed to seal with the agar surface for 15 min, before inverting and incubating at 37°C for 24 hours. The zone of inhibition (ZOI) was measured.

### *In vitro* biofilm disruption assay and confocal microscopy

The overnight culture of PAO1-GFP was diluted in tryptic soy broth (TSB) to an OD600 of 0.05 and kept on ice. In each well of a glass bottom 24-well plate (Cellvis, USA, Cat. No. P24-1.5H-N), 250 µL of top agar was first allowed to solidify after which 300 µL of inoculum was added. The inoculum was permitted to adhere for 30 min at 37°C, after which the media was gently replaced with 500 µL of fresh TSB. After 5 hours, each well was replaced with 800 µL of fresh TSB and was then incubated overnight. At 24 hours, wells were gently rinsed with 800 µL of PBS+Mg, then replenished with 300 µL of modified-LB. Treatments were subsequently applied as followed: control treatment (untreated) consisted of 200 µL of PBS+Mg; phage treatment (PAML-31-1) consisted of 200 µL of phage suspension at 10^11^ PFU/mL added directly into the well; control gel treatment was 100 µL of HA-PEG gel in transwell insert topped with 200 µL of PBS+Mg; and HydroPhage (HP) treatment was 100 µL of HA-PEG gel containing PAML-31-1 at 10^11^ PFU/mL in a transwell insert topped with 200 µL of PBS+Mg. Biofilms were monitored using a Leica SP8 laser-scanning confocal microscope with a 25x 0.95-NA water immersion objective at 0, 3, 8, and 21 hours. Biofilm base density was quantified as the percentage of area exhibiting PAO1-GFP signal at the interface between the biofilm and the agar. The change in biofilm density change was calculated at 21 hours relative to the initial density. Given the significant presence of planktonic bacteria and the disruption induced by treatments, biofilm thickness was manually determined from the media–biofilm boundary to the base of the biofilm using x–z and y–z projections. Three-dimensional reconstructions were generated using Imaris imaging software (version 9.9, Oxford Instruments).

An extensively drug-resistant (XDR) clinical *P. aeruginosa* isolate (CPA012), resistant to amikacin, aztreonam, ceftazidime, ciprofloxacin, doripenem, imipenem, levofloxacin, meropenem, and tobramycin, with intermediate resistance to piperacillin and tazobactam, was also evaluated. CPA012 biofilms were established over 48 hours with four TSB medium changes. Following biofilm formation, wells were treated with either control, PAML-31-1, or HP for 17 hours. Biofilms were then gently rinsed with saline and stained with BacLight (Invitrogen, USA, Cat. No. L7012) for 15 minutes at room temperature. The BacLight staining solution was prepared by mixing SYTO9 and propidium iodide in a 1:1 ratio, using 10 µL of the dye mixture per 1 mL saline. Biofilm base density, accounting for both live and dead cells, was normalized to that of the untreated control group.

### *In vitro* cytotoxicity and activation of hydrogel against RAW264.7 macrophages

A transwell exposure assay was used to measure the cytotoxicity and immunogenicity of pre-formed hydrogels against RAW264.7 murine macrophage-like cells **(Fig. 5c)**. RAW264.7 cells were grown to passage 6 post-thaw in complete DMEM (DMEM/F12 + 10% fetal bovine serum + 1% penicillin/streptomycin). 5 x 10^4^ cells were seeded in 24-well tissue culture-treated plates. After overnight incubation, the media was replaced with 500 μL of fresh cDMEM. Cells were treated for 24 hours with one of the following: media only (negative control), 8% DMSO (positive cytotoxicity control), 1 μg/mL *Escherichia coli* lipopolysaccharide (M1 polarizing positive control; Sigma-Aldrich, Cat. No. L3024-5mg), 20 ng/mL mouse IL-4 (M2 polarizing positive control; ThermoFisher Scientific, USA, Cat. No. 214-14-1MG), FPLC-purified PAML-31-1 at 5 x 10^9^ PFU/mL, or 100 μL of pre-formed hydrogels in transwell inserts, with 200 μL of fresh media added on top to prevent drying. After 24 hours of incubation, the media was removed, and cells were rinsed once with PBS. Cells were then stained with 300 μL of 1:3000 IRDye 800CW in PBS for 15 min at room temperature. Staining was quenched by the addition of 1 mL ice-cold FACS buffer. After centrifugation, cells were resuspended in 300 μL of anti-mouse CD16/32 (Fc block) in FACS at a concentration of 1 μg/million cell for 10 min at 4°C. After centrifugation and removal of excess Fc block, samples were stained with antibodies **(Extended Data Table 2)** for 20 min at 4°C in a staining volume of 300 μL. Samples were rinsed twice in ice-cold FACS and fixed in 500 μL 10% neutral buffered formalin (ThermoFisher Scientific, USA, Cat. No. 9990245) for 15 min at room temperature. Cells were scraped and transferred to a 96-well V-bottom plate for analysis by flow cytometry within one week.

### *In vitro* and *in vivo* bioluminescence to CFU (BLI-CFU) correlation

Quantification of bacterial wound infections using luminescent strains of *P. aeruginosa* was performed as described previously^31^. In brief, a dilution series of *Pseudomonas aeruginosa* PAO1-Xen41 (PerkinElmer, MA, USA) was prepared in a black 96-well plate (ThermoFisher Scientific, USA, Cat. No. 165305) ranging from 10^0^ to 10^7^ CFU/well in a total volume of 100 μL. Samples were prepared in triplicate. CFU counts were verified by plating serial dilutions on nonselective LB agar plates. Bioluminescence imaging was performed on the Lago X *In Vivo* Imaging System (Spectral Instruments Imaging, USA), using the same settings described in the Methods section on “Bioluminescence Imaging”. Linearity between bioluminescent flux (log_10_(photons/s)) and bacterial burden (CFU) were assessed for flux values above 4 and 5 for the *in vitro* and *in vivo* experiments, respectively.

### Delayed treatment mouse infection model

Male C57BL/6J mice (#000664) were purchased from the Jackson Laboratory. All experiments and animal use procedures were approved by the Institutional Animal Care and Use Committee (IACUC) at the School of Medicine at Stanford University. The study design was adapted from previous work^72^. Briefly, 6-7-week-old male mice were anesthetized using 3% isoflurane and shaved. The shaved area was cleaned with povidone-iodine (Dynarex, USA, Cat. No. 1201) and isopropyl alcohol swabs. Mice received a single subcutaneous injection of slow-release buprenorphine (0.1-0.5 mg/kg, ZooPharm, USA) as an analgesic. A single circular midline dorsal full-thickness wound was created using a 6 mm biopsy punch (Integra Miltex, Catalog No.12-460-412). The wounds were immediately covered with Tegaderm (3M, USA, Cat. No. 1642W) and secured with silicone adhesive on the periphery (Vapon, USA, Cat. No. NTL1). Mice were placed in padded plastic collars (Saf-T-Shield, KVP International, USA, Cat. No. RCM) to prevent the removal of wound dressings. Mice received 500 μL of 0.9% normal saline by IP injection on the wounding day and food pellets covered with veterinary nutrient supplements (California Veterinary Supply, USA, Cat. No. HS-10033).

On the infection day, *P. aeruginosa* PAO1-Xen41 was grown in LB at 37°C with shaking to mid-log phase. Bacteria were pelleted by centrifugation, resuspended in PBS, and diluted to 1.25 x 10^7^ CFU/mL. The inoculum density was verified by plating serial dilutions on LB agar plates. 24 hours post-wounding, wounds were inoculated with 40 μL PAO1-Xen41 under Tegaderm using an insulin syringe (28G) at 5 x 10^5^ CFU/wound, chosen as the inoculum because it resulted in a 100% infection rate. Infection rate was defined as the percentage of PBS-treated control mice that had CFU counts at the endpoint of the study, Day 6-post infection. Bioluminescence images were obtained 1 hour post-infection using the Lago X *In Vivo* Imaging System to confirm that the inoculum was properly injected under the Tegaderm.

24 hours post-infection (“Day 0”), all mice were imaged before the first treatment. Mice were treated with 1) PBS as control; 2) IV phage via dorsal penile vein injection, 1 x 10^11^ PFU/mL, 40 µL, 4 x 10^9^ PFU total; 3) daily IP injection for 4 days (IP*4), 2.5 x 10^9^ PFU/mL, 400 µL/dose, 4 x 10^9^ PFU total; 4) topical single high-dose (HD), 1 x 10^11^ PFU/mL, 40 µL, 4 x 10^9^ PFU total; 5) topical single low-dose (LD), 2.5 x 10^10^ PFU/mL, 40 µL, 1 x 10^9^ PFU total; 6) topical daily low-dose (DD) for 4 days, 2.5 x 10^10^ PFU/mL, 40 µL/dose, 4 x 10^9^ PFU total; or 7) topical HydroPhage (HP), 1 x 10^11^ PFU/mL, 40 µL, 4 x 10^9^ PFU total. All mice were weighed daily. Additional Tegaderm dressings were applied on top of existing ones if the dressing became loose. Mice that completely removed the Tegaderm dressings or which partially removed and exhibited a resultant >2-log reduction in luminescent flux in a day, indicating potential drying of infection or treatment, were excluded from subsequent analysis. Six days post-infection, mice were euthanized, and the wound was harvested (1 cm^2^). Each wound was bisected. One half was weighed, homogenized, and used for enumerating colony-forming units (CFU) and plaque-forming units (PFU), while the other half was processed for flow cytometry. The LoD of CFU and PFU were defined as 100 divided by the max wound tissue weight in mg of that experiment. The factor of 100 was derived from the theriotical 100 CFU or PFU in 1 mL of sample when 1 colony or plaque was observed on a 10 µL drop of sample on agar. We plotted the highest and lowest LoD of independent experiments on all CFU and PFU plots. The next day, at least 10 colonies from each wound isolate were picked, cultured overnight, and then stored in 30% glycerol at −80°C for further characterization, including quantification of phage resistance using PSA.

### Bioluminescence imaging (BLI)

Bioluminescence imaging was performed on the Lago X *In Vivo* Imaging System (Spectral Instruments Imaging, USA). A series of exposures ranging from 5 s to 60 s were taken to determine the optimal exposure time. The longest exposure without overexposure was used for analysis. The following imaging settings were used: F-stop 1.2, Binning: Heavy, FOV: 25. Images were analyzed using Aura version 4.0 (Spectral Instruments Imaging, USA). Radiance (photons/s/cm^2^/sr) within a manually selected region of interest (ROI) was integrated to measure total signal flux (photons/s).

### Wound flow cytometry

Wounds were rinsed once in HBSS, blotted to remove excess HBSS, and placed into pre-weighed 1.5 mL polystyrene tubes containing 500 μL freshly prepared, ice-cold digestion buffer (RPMI-1640 + 25 mM HEPES, pH 7.4, 1.3 WU/mL Liberase TL (Roche, Switzerland, Cat. No. 05401020001), 2 U/mL bovine DNase I (Roche, Switzerland, Cat. No. 11284932001), 25 mM MgCl_2_, 1 mM CaCl_2_, 200 U/mL collagenase IV (Worthington Biochemical, USA. Cat. No. LS004188)^73^. Samples were weighed and minced finely using scissors. An additional 500 μL ice-cold digestion buffer was added to samples and incubated at 37°C with shaking at 750 x *rpm* for 2 hours. Samples were inverted several times every 30 min during the incubation. After the 2-hour digestion, samples were treated with 100 μL Accutase (Innovative Cell Technologies, San Diego, CA, USA, Cat. No. AT104) and incubated for 20 min at room temperature.

Digested samples were passed through a 40 μm nylon cell strainer (Corning, Corning, NY, USA, Cat. No. 352340) pre-wetted with 1 mL of ice-cold FACS buffer (PBS + 1 mM EDTA + 1% BSA). Samples were gently pressed through the strainer using the rubber end of a syringe plunger. Cell strainers were rinsed with 5 mL of ice-cold FACS buffer. Cells were pelleted by centrifugation (500 x *g*, 10 min, 4°C), and the supernatant was aspirated. Pellets were resuspended in 1 mL ice-cold FACS buffer and pelleted again. Pellets were resuspended in 250 μL of room temperature ACK red blood cell lysis buffer (0.15 M NH_4_Cl, 10 mM KHCO_3_, 0.1 mM EDTA, pH 7.3) and incubated for precisely 4 minutes (variation <5 seconds), after which lysis was stopped by the addition of 750 μL of PBS.

Cells were then pelleted by centrifugation (500 x *g*, 20 min), removed supernatant, and resuspended in 200 μL ice-cold FACS buffer. 10 μL samples were diluted into a final volume of 200 μL and counted on a flow cytometer. 650,000 cells/sample were aliquoted to a 96-well V-bottom tissue culture-treated plate (Corning, Corning, USA, Cat. No. 07-200-96). The remaining cells were pooled and aliquoted for unstained, single-stained, and fluorescence-minus-one (FMO) controls.

For staining, cells were rinsed once in ice-cold FACS buffer and then stained in 50 μL 1:3000 IRDye 800CW Live/Dead NIR stain in PBS for 15 minutes at room temperature. Staining was quenched by the addition of 150 μL of ice-cold FACS buffer. After centrifugation, cells were resuspended in 30 μL of anti-mouse CD16/32 (Fc block) in FACS at a concentration of 1 μg/million cells for 10 min at 4°C. After centrifugation and removal of excess Fc block, samples were stained with antibodies (listed below) for 20 min at 4°C in a staining volume of 30 μL. Samples were then rinsed twice in ice-cold FACS and fixed in 200 μL 4% paraformaldehyde (20 min, room temperature). Samples were then resuspended in 200 μL FACS and acquired on the Cytek Aurora NL-3000 spectral flow cytometer (Cytek Biosciences, CA, USA) using SpectroFlo 3.0 software within 1 week of fixation. Spectral unmixing was performed using narrow gates and without unmixing autofluorescence.

### Whole genome sequencing

Overnight cultures of *P. aeruginosa* PAO1-Xen41 and selected isolates from HP-treated mice were grown in 5 mL of LB medium. These cultures were then subcultured at a 1:25 dilution for 3 hours and subsequently adjusted to an OD600 of 1. A total of 12 mL of bacterial suspension was collected in a conical tube and centrifuged at 2,000 × *g* for 10 minutes. The supernatant was discarded, and the pellet was weighed (30∼50 mg). After resuspending the pellet in 1 mL of PBS and centrifuging again, the supernatant was removed. Finally, the pellet was resuspended in 0.5 mL of 1× DNA/RNA Shield reagent (Zymo Research, USA, Cat. No. R1100). Samples were then sent to Plasmidsaurus (USA) for DNA extraction and whole genome sequencing.

The resulting sequences were analyzed using Geneious Prime (version 2025.0.2, Biomatters Ltd). The wild-type PAO1-Xen41 reference sequence was aligned with the isolates sequences using the LASTZ alignment tool integrated within Geneious. Default alignment parameters were applied, with adjustments for sensitivity and specificity as needed. Single nucleotide polymorphisms (SNPs) were identified and visualized.

### Statistical analyses

Statistical analyses were performed using GraphPad Prism version 10. Ordinary one-way analysis of variance (ANOVA) with Tukey post-hoc tests was used for multiple comparisons unless otherwise noted. Unpaired two-tailed Student’s *t*-tests were used for comparison between two groups unless otherwise mentioned. *P* ≤ 0.05 was considered significant unless otherwise noted. Data were presented as mean with error bars representing the standard deviation (S.D.), unless otherwise noted. (* *P* ≤ 0.05, ** *P* ≤ 0.01, *** *P* ≤ 0.001, **** *P* ≤ 0.0001).

## Acknowledgements

*In vivo* imaging was performed at Stanford Center for Innovation in *In Vivo* Imaging (Sci^3^). We thank Drs. A. Khosravi, N. Nagy, A. Saraswathibhatla, and D. Indana and other members from the Bollyky and Chaudhuri labs for their scientific inputs; Dr. E. Burgener for providing the clinical *Pa* isolate; the staff at the Sci^3^ for providing the support needed to perform the required experiments, including Dr. F. Habte. The research reported in this publication was supported by a Stanford Catalyst Grant and a Stanford Coulter Translational Grant. Q. C. discloses support from Cystic Fibrosis Foundation CHEN21F0, Cystic Fibrosis Research Institute, and Stanford Maternal & Child Health Research Institute. T.D. discloses support from Stanford University Medical Scientist Training Program National Institutes of Health grant T32-GM007365; Stanford Interdisciplinary Graduate Fellowship affiliated with Sarafan ChEM-H, Gold Family Graduate Fellow. M.H. discloses support from National Institutes of Health grants R00-EB028838 and K99EB028838. O.C. acknowledges support from NIH grants R01 AR081993 and R01 GM148535. P.L.B discloses support from National Institutes of Health grant R01 HL148184-01; National Institutes of Health grant R01 AI12492093; National Institutes of Health grant R01 DC019965; Cystic Fibrosis Foundation grant; Emerson Collective grant. The contents are those of the authors and do not necessarily represent the views of the funding agencies.

## Author contributions

Y.H.L., T.D., and Q.C. contributed equally. Y.H.L., T.D., Q.C., R.M., D.A., O.C., and P.L.B. conceived the study. Y.H.L. and R.M. performed hydrogel phage release pilot studies. Y.H.L., T.D., and Q.C. designed the experiments and mouse model. Y.H.L. and L.J.Z. performed rheological tests, and Y.H.L. conducted hydrogel degradation and bead diffusion studies. Y.H.L., T.D., Q.C., L.J.Z., A.E., C.D.L., and W.S. performed *in vitro* phage release studies. Y.H.L., A.E., Z.L., and T.H.W.C performed *in vitro* phage selection and resistance assays. Y.H.L. and L.J.Z. conducted biofilm studies. T.D., N.A.P., and H.A.M. performed the flow cytometry assays and *in vitro* macrophage cytotoxicity and activation studies. Q.C., T.D., A.E., Y.H.L., Z.L., L.J.Z., F.G.B., and T.H.W.C. performed *in vivo* studies, with C.D.L, Y.H.L., and A.E. curating data. M.H. conducted TEM imaging. A.E., L.J.Z., T.D., T.H.W.C., Z.L. and A.H. propagated and purified phages. Y.H.L., Q.C., T.D., A.E., and J.P. analyzed and interpreted the data. O.C. and P.L.B. supervised the study. Y.H.L., T.D., Q.C., O.C., and P.L.B. wrote the manuscript. All authors reviewed the manuscript.

## Data availability statement

The data supporting the results in this study are available within the paper and Extended Data. All data acquired during the study are available from the corresponding author on reasonable request.

