## Extended Data for "Dosing and Delivery of Bacteriophage Therapy In a Murine Wound Infection Model"

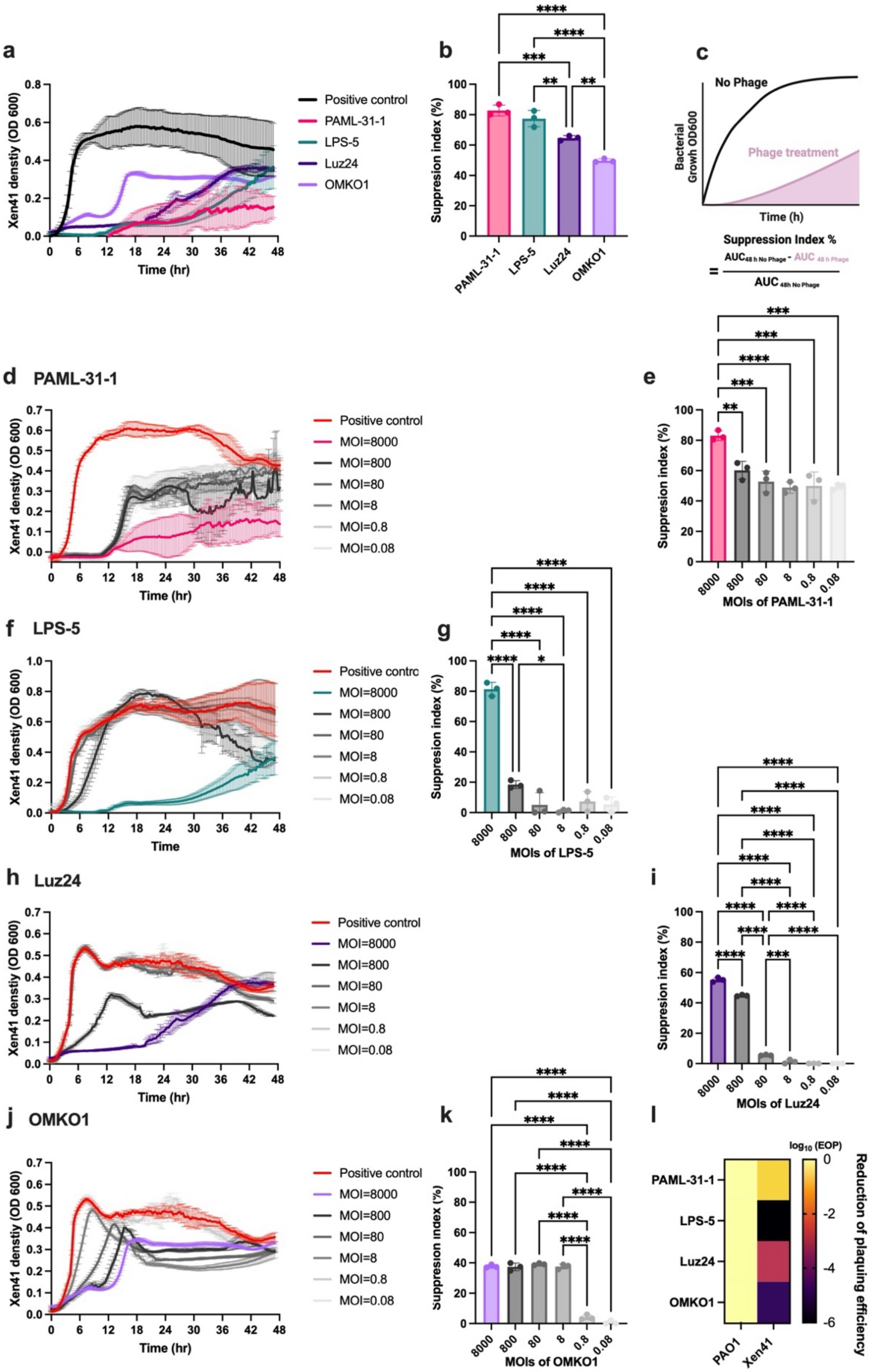

**Extended Data Fig. 1. PAML-31-1 phage is most effective against PAO1-Xen41 in both planktonic and on-agar conditions.** **a**,**b**,**c**, Four different antipseudomonal phages—PAML-31-1, LPS-5, Luz24, and OMKO1—were co-cultured with *P. aeruginosa* strain PAO1-Xen41 in planktonic condition at the multiplicity of infection (MOI) of 8,000 with shaking for 48 hours and monitored every 20 minutes. Time evolution of optical density at 600 nm (OD600) (**a**), the quantification of bacterial suppression by suppression index (SI%) (**b**); n=9 for positive control and n=3 for each phage, and the definition of SI% (**c**). **d**-**k**, Various phages were co-cultured with PAO1-Xen41 bacteria at a range of MOIs. Time evolution of OD600 for PAML-31-1 (**d**), LPS-5 (**f**), Luz24 (**h**), and OMKO1 (**j**) treatments, and the quantification of SI% for PAML-31-1 (**e**), LPS-5 (**g**), Luz24 (**i**), and OMKO1 (**k**) treatments; n=3 for each MOI. **i**, The efficiency of plaquing (EOP) was quantified by comparing the log reduction of plaque counts of different phages on *P. aeruginosa* strain PAO1-Xen41 versus its normalized titer using parent strain PAO1, n=1 each. Data are mean ± SD. One-way ANOVA with Tukey's post-hoc tests for (**b**), (**e**), (**g**), (**i**), (**k**). (* $P \leq 0.05$, ** $P \leq 0.01$, *** $P \leq 0.001$, **** $P \leq 0.0001$).

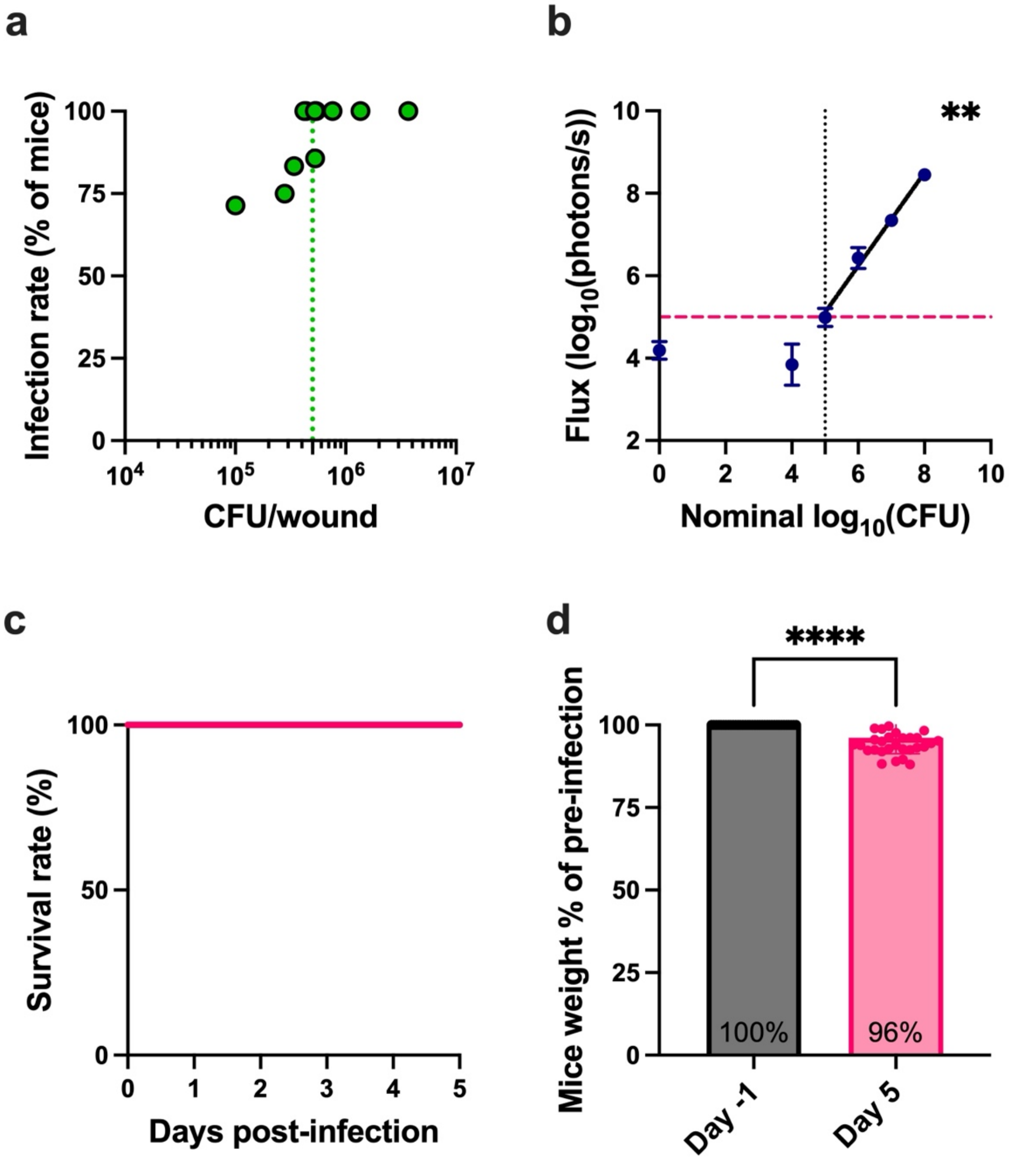


**Extended Data Fig. 2. Murine wound infection model.** **a**, Dose-dependent infection rate following wound inoculation with increasing bacterial colony-forming units (CFU). Green dashed line indicates the inoculum of $5x10^5$ CFU/wound. **b,** bioluminescent signal and nominal CFU correlation above $1x10^5$ CFU for PAO1-Xen41 *in vivo* (Pearson r = 0.9957, P < 0.01; $R^2$ = 0.9768). **c,** A 100% survival rate was observed across the 5-day post-infection period. **d,** Minimal impact on mouse body weight between day -1 and day 5 post-infection, shown as percentage of pre-infection weight. Data are presented as mean ± S.D for (**b**) and (**d**). Linear regression (R = 0.9768) and Pearson r correlation (r = 0.9957) above $1x10^5$ CFU for (**d**). Unpaired two-tailed Student's *t*-test for (**d**). (** *P* ≤ 0.01, **** *P* ≤ 0.0001).

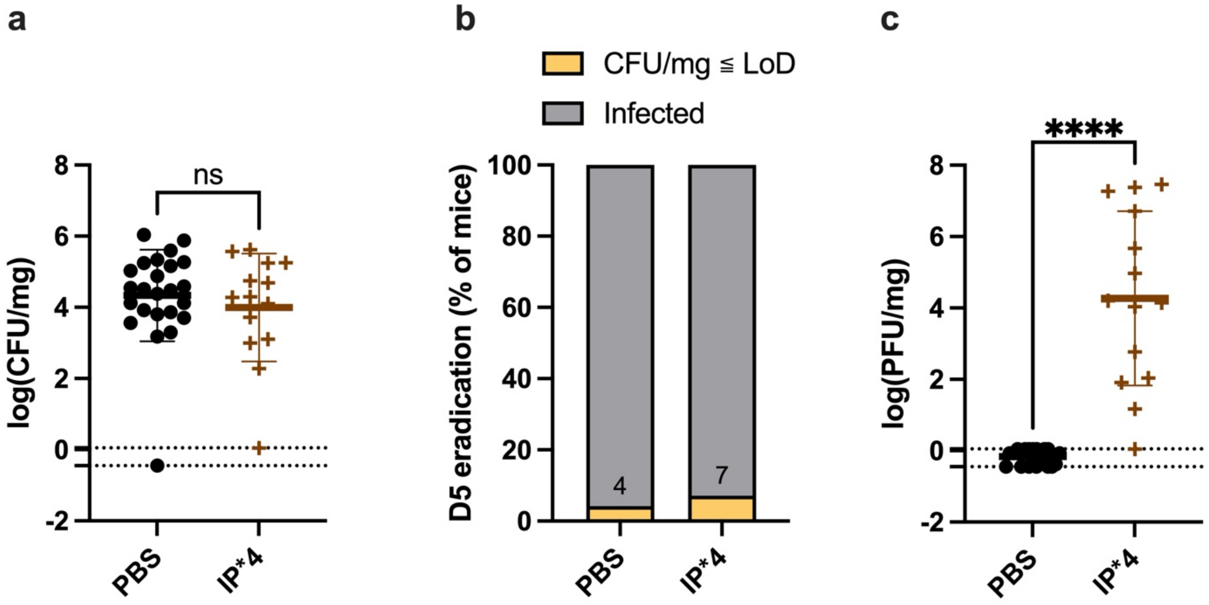


**Extended Data Fig. 3. Repeated intraperitoneal (IP) phage administration fails to eradicate bacterial infection in wounds. a**, Bacterial burden quantified as log(CFU/mg) in wounds, showing no significant difference between PBS control and IPx4 treatment groups (ns = not significant; dots represent LoD). **b,** The eradication rate (CFU-based) achieved by treatments on Day 5 post-treatment. c, the phage counts (PFU/mg) recovered from harvested mice wound tissue. Data are presented as mean ± S.D. Unpaired two-tailed Welch's *t*-test for (**a**) and (**c**). (**** $P \leq 0.0001$).

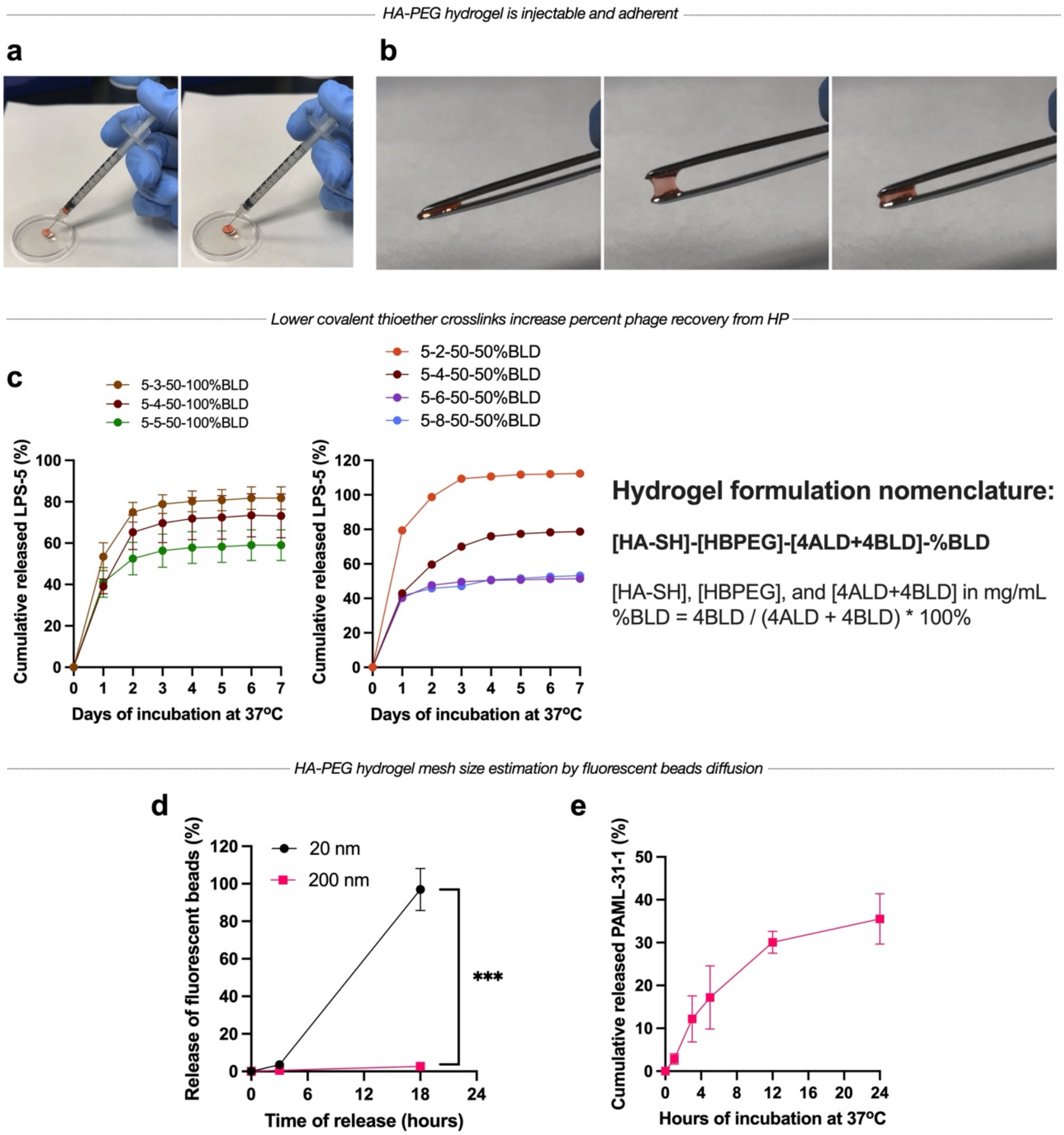


**Extended Data Fig. 4. HA-PEG hydrogel mechanics and particle release. a,** HA-PEG hydrogel was injectable through the fine 28G needle. **b**, HA-PEG hydrogel was sticky as shown by the adherence to both sides of the tweezers. Hydrogel was dyed with phenol red and the formulation was 5 mg/mL HA-SH, 1 mg/mL HBPEG, and 75 mg/mL of 4-arm PEGs with 10% 4BLD and 90% 4ALD. **c**, Lower covalent thioether crosslinks increase percent phage recovery from HP. LPS-5 phage was encapsulated at $10^{11}$ PFU/mL in HP. n = 3 gels each with technical triplicates (left) and n = 1 gel each with technical triplicates (right)**.** The nomenclature of HA-PEG hydrogel formulation was in the order of concentration in mg/mL of polymers—HA-SH, HBPEG, 4-arm PEGs—followed by the percentage of benzaldehyde (%BLD) in 4-arm PEGs. For example, 5-1-75-10%BLD represents 5 mg/mL HA-SH, 1 mg/mL HBPEG, 75 mg/mL of 4-arm PEGs with 10% 4BLD and 90% 4ALD. **d**, the cumulative release curve of 20 and 200 nm fluorescent beads from HA-PEG hydrogels for 18 hrs. n=2 for 20 nm and n=3 for 200 nm beads. **e**, the cumulative release curve of PAML-31-1 phage from HP for 24 hrs. n=3 gels. Data are presented as mean ± S.D for (**c**), (**d**) and (**e**). Two-way ANOVA with Geisser-Greenhouse correction and Tukey's post-hoc tests for (**d**). (*** $P \leq 0.001$).

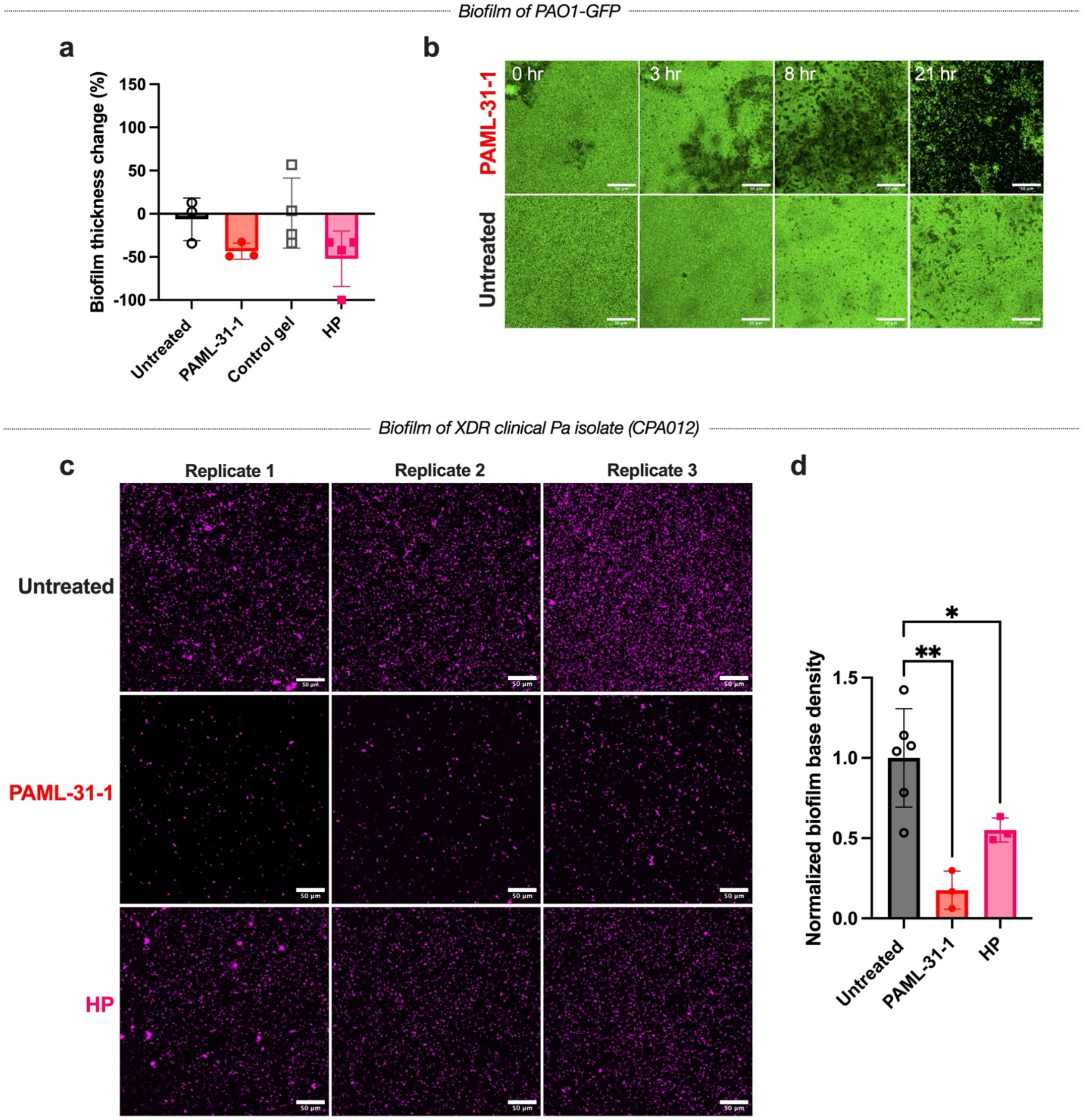


**Extended Data Fig. 5. HP and PAML-31-1 disrupt *P. aeruginosa* biofilm. a**,**b**, Biofilm disruption of PAO1-GFP. The change of thickness as a percentage of initial thickness for each biofilm after exposure to treatment for 21 hours at 37°C (**a**); n=3, 3, 4, and 4 replicates for untreated, PAML-31-1, control gel, and HP groups, respectively; n = 2 independent experiments each; The representative images of biofilm treated with PAML-31-1 in media or untreated control over time (**b**). Scale bar: 50 µm. **c**,**d**, Biofilm disruption of an extensive drug resistant (XDR) clinical *P. aeruginosa* isolate (CPA012). The representative images of biofilm treated with PAML-31-1 in media, HP, or untreated control (**c**) and quantification of biofilm base density normalized to untreated group (**d**). n=6 for untreated control and n=3 for PAML-31-1 and HP treatments. Scale bar: 50 µm. Data are presented as mean ± S.D for (**a**) and (**d**). One-way ANOVA with Tukey's post-hoc tests for (**d**). (* $P \leq 0.05$).

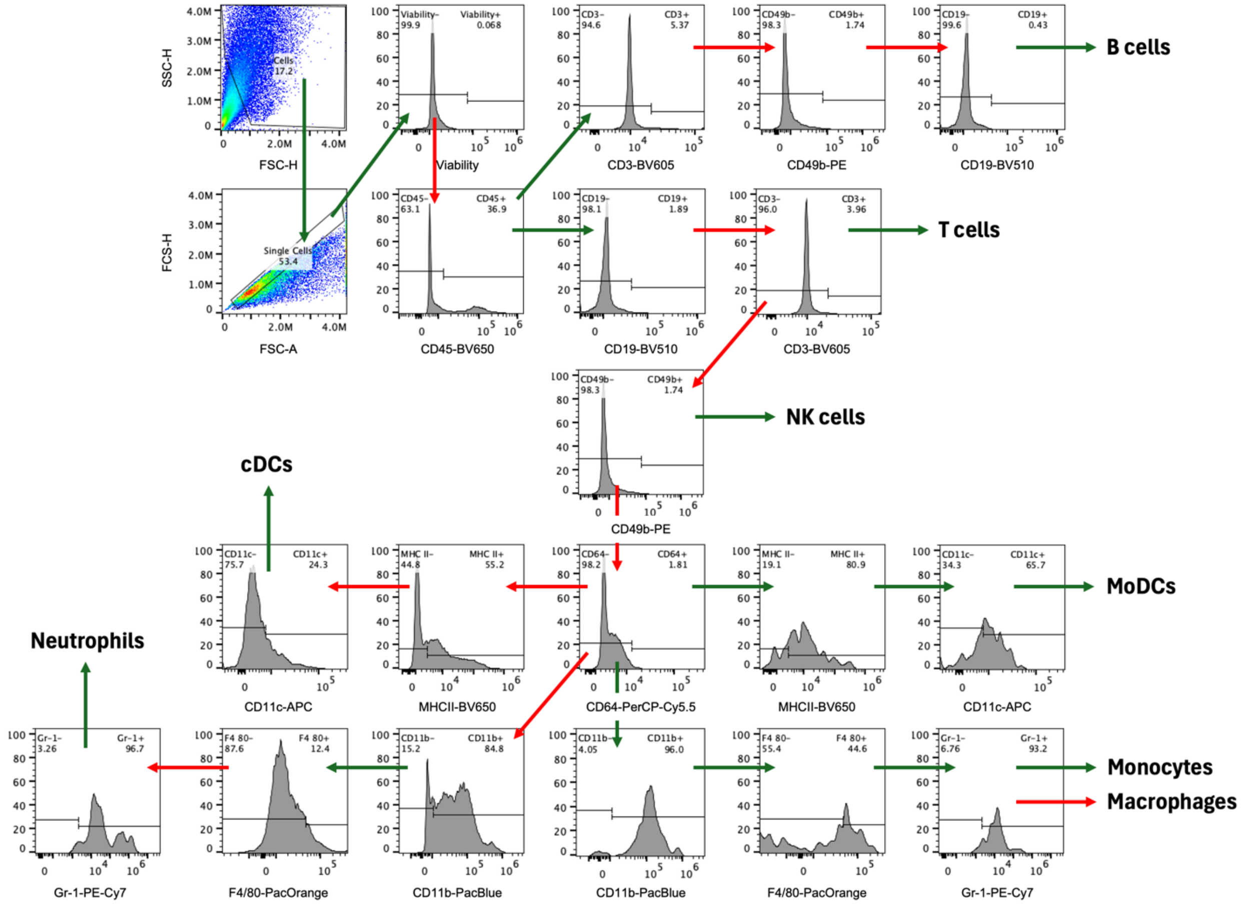


**Extended Data Fig. 6. Gating strategy for wound flow cytometry.** A 13-color panel was used to perform spectral flow cytometry to discriminate major myeloid and adaptive immune cell populations in dissociated wound tissues collected on Day 6 post-infection. Manual binary gating using fluorescence-minus-one (FMO) controls was used to set gates. Modal histograms are shown with y-axis as percentages. Green arrows indicate forward gating of positive populations, while red arrows indicate forward gating of negative populations. The following cell types were identified: CD45$^{+}$ cells, T cells (CD45$^{+}$CD19$^{-}$CD3$^{+}$), B cells (CD45$^{+}$CD3$^{-}$CD49b$^{-}$CD19$^{+}$), NK cells (CD45$^{+}$CD19$^{-}$CD3$^{-}$CD49b$^{+}$), Innate immune cells (LIN$^{-}$; CD45$^{+}$CD19$^{-}$CD3$^{-}$CD49b$^{-}$), conventional dendritic cells (cDCs; LIN$^{-}$CD64$^{-}$MHCII$^{+}$ CD11c$^{+}$), monocyte-derived dendritic cells (MoDCs; LIN$^{-}$CD64$^{+}$MHCII$^{+}$CD11c$^{+}$), neutrophils (LIN$^{-}$CD64$^{-}$CD11b$^{+}$F4/80$^{-}$Gr-1$^{+}$), macrophages (LIN$^{-}$CD64$^{+}$CD11b$^{+}$F4/80$^{+}$Gr-1$^{-}$), and monocytes (LIN$^{-}$CD64$^{+}$CD11b$^{+}$ F4/80$^{+}$Gr-1$^{+}$).

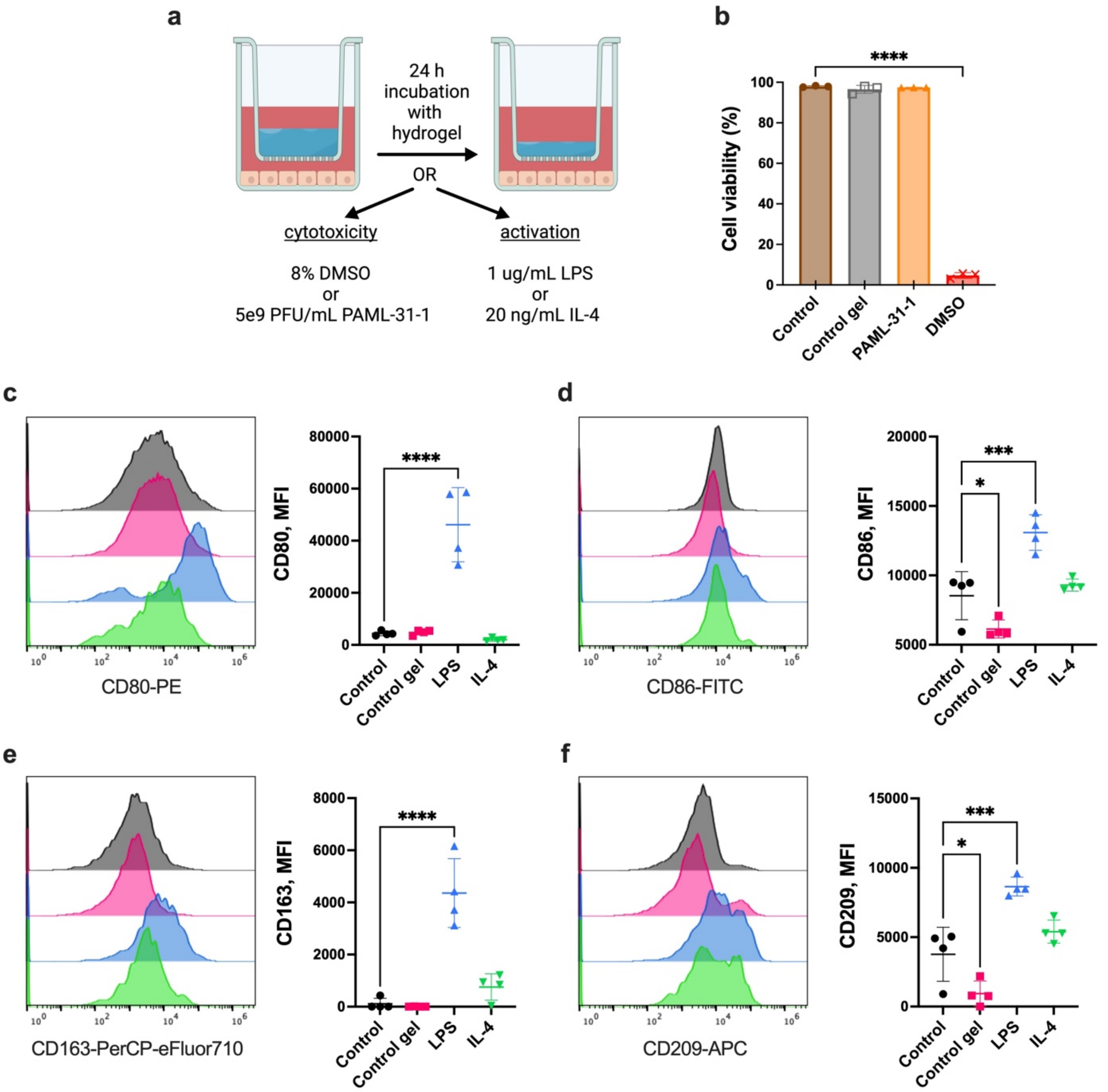


**Extended Data Fig. 7. HA-PEG hydrogel is not cytotoxic or activating towards RAW264.7 macrophages. a,** Schematic of transwell exposure assay of cytotoxicity and macrophage activations. **b**, Cytotoxicity of HA-PEG gel (control gel) and PAML-31-1 phage to RAW264.7 cells using transwell system; n = 3 each. **c-f**, Representative histograms (left) and median fluorescence intensity (MFI) of fluorochrome-conjugated antibodies against classical M1 (CD80, CD86) and M2 (CD163, CD209) markers in RAW264.7 cells after treatment with media only control, control gel in transwell system, 1 ug/mL *E. coli* lipopolysaccharide (LPS), or 20 ng/mL mIL-4; n = 4 each. Data are mean ± S.D. Ordinary one-way ANOVA with Tukey post-hoc tests for (**b**) and with Dunnett post-hoc tests for (**c-f**) (* $P \leq 0.05$, *** $P \leq 0.001$, **** $P \leq 0.0001$).

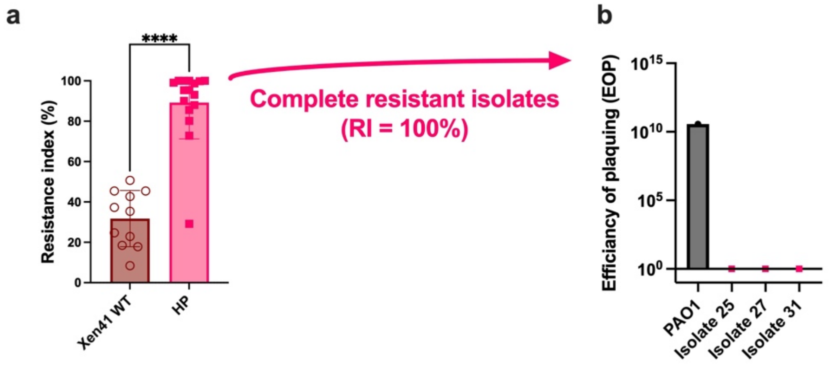


**c**

| Prior treatment | Mice ID | Gene(s) | Role of Gene(s) | Mutation Type | Amino acid Change |
|---|---|---|---|---|---|
| **HydroPhage (HP)** | *022024-#25* | *lolE* | Lipoprotein localization | Substitution | 2: PLPF → PLSV<br>11: L → T<br>20: N → S<br>23: I → V<br>40: A → V<br>43: I → V<br>52: QK → DR<br>65: H → Q<br>68: I → V<br>69: SA → ES<br>71: T → Y<br>73: PL → PI<br>78: QT → RA<br>80: V → L<br>82: N → D<br>257: T → S<br>258: Q → H<br>260: S → N<br>262: F → Y<br>263: N → Q<br>265: MK → IR<br>280: I → V<br>289: A → S<br>292: I → V<br>296: A → T<br>299: RT → KS<br>313: R → G<br>318: I → T<br>332: V → L<br>340: F → V<br>345: ITGMIDRIER → VSAWISALEK<br>356: V → L<br>359: KVFS → QFLA<br>369: N → D<br>374: D → Q<br>376: QVL→ MLD<br>383: I → V<br>385: S → G<br>389: LM → VL<br>393: L → F<br>399: S → A<br>410: S → A |
| | *022024-#27* | *migA* | LPS core biosynthesis | Substitution | 196: T → P |

**Extended Data Fig. 8. Resistance to PAML-31-1 is attributed to mutations in lipopolysaccharide (LPS). a**,**b**, HP treated mice isolates showing complete resistant to PAML-31-1 in PSA (**a**) and EOP (**b**) were selected for whole genome sequencing (WGS). **c**, WGS reveals point mutations in lipoprotein localization and LPS core biosynthesis genes.

**Extended Data Table 1. Antibodies used for flow cytometry on dissociated wound tissues.**

| Antibody | Color | Clone | Host | Isotype | Dilution | Brand | Cat. No. |
|---|---|---|---|---|---|---|---|
| IRDye 800CW | | | | | | Li-COR Biosciences | 929-70020 |
| Fc block | | 93 | Rat | IgG2a lambda | 1 ug/1M cells | BioLegend | 101320 |
| Brilliant Stain buffer | | | | | 1:20 | BD Biosciences | 563794 |
| CD11b | PacBlue | M1/70 | Rat | IgG2b kappa | 1:100 | BioLegend | 101224 |
| CD19 | BV510 | 6D5 | Rat | IgG2a kappa | 1:20 | BioLegend | 115546 |
| F4/80 | PacOrange | BM8 | Rat | IgG2a | 1:20 | Invitrogen | MF48030 |
| CD3 | BV605 | 17A2 | Rat | IgG2b kappa | 1:40 | BioLegend | 100237 |
| CD45 | BV650 | 30-F11 | Rat | IgG2b kappa | 1:100 | BioLegend | 103151 |
| I-A/I-E | FITC | M5/114.15.2 | Rat | IgG2b kappa | 1:100 | BioLegend | 107643 |
| CD64 | PerCP-Cy5.5 | X54-5/7.1 | Mouse | IgG1 kappa | 1:50 | BioLegend | 139307 |
| CD49b | PE | HMα2 | Hamster | IgG | 1:100 | BioLegend | 103506 |
| Gr-1 | PE-Cy7 | RB6-8C5 | Rat | IgG2b, kappa | 1:100 | BioLegend | 108416 |
| CD11c | APC | N418 | Hamster | IgG | 1:50 | BioLegend | 117317 |

**Extended Data Table 2. Antibodies used for flow cytometry to assess cytotoxicity and activation of RAW264.7 cells.**

| Antibody | Color | Clone | Host | Isotype | Dilution | Brand | Cat. No. |
|---|---|---|---|---|---|---|---|
| IRDye 800CW | | | | | | Li-COR Biosciences | 929-70020 |
| Fc block | | 93 | Rat | IgG2a lambda | 1 ug/1M cells | BioLegend | 101320 |
| Brilliant Stain buffer | | | | | 1:20 | BD Biosciences | 563794 |
| I-A/I-E | BV510 | M5/114.15.2 | Rat | IgG2b, kappa | 1:100 | BioLegend | 107635 |
| CD86 | FITC | GL-1 | Rat | IgG2a, kappa | 1:100 | BioLegend | 105006 |
| CD163 | PerCP-eFluor710 | TNKUPJ | Rat | IgG2a, kappa | 1:100 | eBioscience | 46-1631-82 |
| CD80 | PE | 16-10A1 | Hamster | IgG | 1:100 | BioLegend | 104708 |
| CD209 | APC | MMD3 | Mouse | IgG2c, kappa | 1:100 | BioLegend | 833006 |
